# Mushroom Body Homology and Divergence across Pancrustacea

**DOI:** 10.1101/856872

**Authors:** Nicholas J. Strausfeld, Gabriella H. Wolff, Marcel E. Sayre

**Affiliations:** Department of Neuroscience, School of Mind, Brain and Behavior, University of Arizona, Tucson, Arizona, USA; Department of Biology, University of Washington, Seattle, Washington, USA; Lund Vision Group, Department of Biology, Lund University, Lund, Sweden

**Keywords:** Learning and memory, Ground pattern organization, Brain Evolution, Crustacea, Insecta

## Abstract

Descriptions of crustacean brains have mainly focused on three highly derived lineages: the reptantian infraorders represented by spiny lobsters, lobsters, and crayfish. Those descriptions advocate the view that dome- or cap-like neuropils, referred to as “hemiellipsoid bodies,” are the ground pattern organization of centers that are comparable to insect mushroom bodies in processing olfactory information. Here we challenge the doctrine that hemiellipsoid bodies are a derived trait of crustaceans, whereas mushroom bodies are a derived trait of hexapods. We demonstrate that mushroom bodies typify lineages that arose before Reptantia and exist in Reptantia. We show that evolved variations of the mushroom body ground pattern are, in some lineages, defined by extreme diminution or loss and, in others, by the incorporation of mushroom body circuits into lobeless centers. Such transformations are ascribed to modifications of the columnar organization of mushroom body lobes that, as shown in *Drosophila* and other hexapods, contain networks essential for learning and memory. We propose that lobed mushroom bodies distinguish crustaceans that negotiate the multidimensionality of complex ecologies, where continuous updating of multistimulus valence and memory is paramount.

## INTRODUCTION

As demonstrated in *Drosophila melanogaster*, paired mushroom bodies in the insect brain play manifold roles in learning and memory (Aso et al., 2014a, b; Cognigni et al., 2018). One diagnostic tool for identifying putative mushroom body homologues in other mandibulates is an antibody raised against the catalytic subunit of protein kinase A, encoded by the *Drosophila* gene DC0 (Kalderon and Rubin, 1988), and required for effective learning and memory (Skoulakis et al., 1993). The antibody, known as “anti-DC0”, selectively identifies the columnar neuropils of mushroom bodies of insects (Farris and Strausfeld, 2003) and of other arthropods with, until very recently, the notable exception of crustaceans (Wolff and Strausfeld, 2015). Indeed, numerous studies have disputed or expressed ambivalence about centers in comparable locations in the crustacean brain being mushroom body homologues (e.g., Strausfeld et al., 1998; Fanenbruck et al., 2004; Fanenbruck and Harzsch, 2005; Farris, 2013; Sandeman et al., 2014; Krieger et al., 2015; Harzsch and Krieger, 2018; Machon et al., 2019; Wittfoth et al., 2019).

The discovery of mushroom bodies in mantis shrimps by Wolff et al. (2017) caused us to revise our view of mushroom bodies in crustaceans and to look for these centers in other lineages of this taxon. Here, we provide evidence that mushroom bodies generally occur in Crustacea, thereby further supporting Hexapoda + Crustacea as the established subphylum Pancrustacea. To arrive at this conclusion, we have considered the following. Do mushroom bodies, defined by their columnar lobes, occur broadly across crustaceans? Are anti-DC0-immunoreactive centers that lack columnar lobes phenotypic homologues of mushroom bodies? Can the reduction or the absence of such centers in certain crustacean lineages be explained by an evolved loss of their ancestral ground pattern?

Historically, identification of the paired insect mushroom body originally relied on just four neuroanatomical traits: an overall fungiform shape; a location exclusively in the brain’s rostrolateral protocerebrum; an internal composition comprising hundreds to tens of thousands of intrinsic neurons originating from minute cell bodies clustered at the brain’s rostral surface; and the possession of columnar lobes from which originate output neurons. An often-included fifth character typical of almost all winged insects is a cap of neuropil called the calyx, comprising the dendrites of intrinsic neurons, which receives inputs from the antennal lobes and other sensory neuropils. Even flightless Zygentoma (silverfish and firebrats) possess calyces (Farris, 2005a), as do Diplura and Collembola, two sister groups of Insecta (Böhm et al., 2012; Kollmann et al., 2011). However, Ephemeroptera (mayflies) and Odonata (dragonflies, darters) are calyxless; inputs to their mushroom bodies supply their columnar lobes directly (Strausfeld et al., 2009). Such distinctions demonstrate a general property of all well-defined protocerebral brain centers: retention of ancestral ground patterns despite evolved modifications such as losses of some components and elaborations of others. For the mushroom body these modifications include: variations of columnar lobe organization; expanded representations of sensory modalities; and hypertrophy of the calyces (Strausfeld et al., 2009).

Despite such variations, the insect mushroom body has additional features that are always present. Foremost is that the columnar lobes are partitioned along their length such that inputs to the lobes are constrained to discrete synaptic domains (Li and Strausfeld, 1997; Ito et al., 1998; Strausfeld 2002). These domains receive functionally defined inputs and supply functionally defined mushroom body output neurons (MBONs; Aso et al., 2014a) that extend to circumscribed regions of the brain’s medial and rostral protocerebrum. There, they contribute to further higher processing and have arrangements comparable to those relating the mammalian hippocampus to cortical areas (Bienkowski et al., 2018). Certain outputs from the columns also project recurrently to more distal levels of the mushroom body itself (Gronenberg, 1987; Zwaka et al., 2018). These arrangements are defined by neurons usually expressing GABA (Liu and Davis, 2009). The same organization typifies the columnar lobes of the stomatopod mushroom bodies where input and output neurons are arranged along the length of the lobes (Wolff et al., 2017). Also apparent across insect species is that subsets of intrinsic neurons parse the mushroom body lobes into longitudinal subunits that are differentiated from each other with regard to their clonal lineages (Yang et al., 1995; Crittenden et al., 1998; Tanaka et al., 2008), as well as by their aminergic and peptidergic identities (Sjöholm et al., 2006; Strausfeld et al., 2000, 2003). In stomatopods, longitudinal divisions are represented by four columnar lobes extending together approximately in parallel (Wolff et al., 2017). In *Drosophila* and other insects these characteristic attributes have been shown to be critical in supporting functions relating to the modality, valence, stored memories and behavioral relevance of information computed by the mushroom body (Li and Strausfeld, 1999; Aso et al., 2014a,b; Owald et al., 2015; Takemura et al., 2017; Hattori et al., 2017; Cognigni et al., 2018). In total, thirteen characters have been identified that define both the insect (*Drosophila*) and mantis shrimp (Stomatopod) mushroom bodies (see, Wolff et al., 2017). We show here that crown eumalacostracan species belonging to lineages originating early in evolutionary history share the same expanded set of diagnostic characters that define insect mushroom bodies, including the partitioning of columns into discrete circuit-defined domains. These can occur not only as segment-like partitions of the columnar lobes but also as tuberous outswellings, as they do in basal Pterygota (Farris, 2005a,b; Strausfeld et al., 2009). A comparable arrangement defines the columnar lobes of the mushroom bodies in the cephalocarid *Hutchinsoniella macracantha*, an allotriocarid basal to Branchiopoda, Hexapoda and Remipedia (Stegner and Richter, 2011; Schwentner et al., 2017).

Because stomatopods are unique amongst crustaceans in possessing elaborated optic lobes that serve a multispectral color and polarization photoreceptor system (Thoen et al., 2017a, 2018), it could be argued that mushroom-bodies in the mantis shrimp are unique apomorphies that have evolved specifically to serve those modalities. However, Stomatopoda are an outgroup of Eucarida (Euphausiacea + Decapoda) and the status of mushroom bodies as the ancestral ground pattern is supported by corresponding centers in the lateral protocerebrum of later evolving eumalacostracan lineages. *Lebbeus groenlandicus*, a member of the caridid family Thoridae, has been shown to possess paired mushroom bodies (Sayre and Strausfeld, 2019). Studies of Reptantia show that at least one anomuran group has mushroom bodies equipped with columnar lobes (Strausfeld and Sayre, 2019). Here we describe neuroanatomical characters defining mushroom bodies also in cleaner shrimps (Stenopodidae) and several groups of carideans.

Many types of reptantian crustaceans, such as crayfish and lobsters, possess domed neuropils lacking columnar lobes (Blaustein et al., 1988). These neuropils were first termed hemiellipsoid bodies (corpo emielissoidale) by Bellonci (1882) as the homologue of the calyx of the insect mushroom body, the columnar lobe being identified as the corpo allungato (Bellonci, 1882). Hanström followed this terminology, using “hemiellipsoid body” as a synonym for centers he considered to be homologues of the insect mushroom body calyx (Hanström, 1925, 1931). Ignoring this nomenclature’s original intent, over the last two decades numerous studies focusing on Reptantia, a relatively recent lineage, insist it is the hemiellipsoid body not the mushroom body that represents the crustacean ground pattern (Kenning et al., 2013; Sandeman et al., 2014; Machon et al., 2019).

Apart from a fundamental misunderstanding of the nomenclature, that viewpoint is incorrect and is in conflict with the demonstration here, and in two recent studies (Wolff et al., 2017; Sayre and Strausfeld, 2019), that mushroom bodies hallmark eumalacostracan lineages that diverged earlier than Reptantia. As will be discussed later, comparisons across Eumalacostraca described here indicate that contrary to received opinion mushroom bodies are ubiquitous across Crustacea. In contrast to the conserved mushroom body ground pattern in insects, evolved modifications of the mushroom body ground pattern in crustaceans have resulted in highly divergent morphologies, including centers lacking defined lobes, both within Caridea and in certain lineages of Reptantia.

## MATERIALS AND METHODS

To screen for putative mushroom bodies or centers in corresponding locations we employed anti-DC0 immunostaining of lateral protocerebra. Identified centers were then subjected to various histological techniques to detect characteristic arrangements that define the hexapod-stomatopod mushroom body ground pattern. The nuclear stain syto13 was used to identify neuronal cell bodies and to distinguish minute basophilic cell bodies associated with anti-DC0-immunoreactive centers. Immunohistology was used to reveal neuroarchitectures as well as putative serotoninergic and dopaminergic neurons belonging to mushroom body input or output neurons (MBINs, MBONs; Aso et al., 2014a,b) and feedforward or recurrent GABAergic elements within mushroom body circuitry (Perisse et al., 2016; Bicker et al., 1985; Leitch and Laurent, 1996). Reduced silver staining and *α*-tubulin immunohistology was employed to reveal the fibroarchitecture of the lateral protocerebrum. Golgi impregnation methods revealing quantities of single neurons were used to compare neuronal organization in mushroom bodies and their suspected “hemiellipsoid body” homologues in Reptantia.

### Animal collection

Animals used in this study were obtained from a variety of commercial vendors. Coral banded cleaner shrimps (*Stenopus hispidus* n=25) and fiddler crabs (*Uca minax* n=12) were purchased from Salty Bottom Reef Co. (New Port Richey, FL). Land hermit crabs (*Coenobita clypeatus* n=13), the crayfish *Orconectes immunis* (n=8) and *Procambarus clarkii* (Girard, 1852 n=10), and cockroaches (*Periplaneta americana* n=20) were obtained from Carolina Biological Supply Co. (Burlingham, NC). Pistol shrimps (*Alpheus bellulus* n=20) and some *S. hispidus* (n=3) were purchased from LiveAquaria (Rhinelander, WI). White-legged shrimp *Penaeus vannamei* (n=24) were purchased from Gulf Specimen Marine Laboratories (Panacea, FL). Stomatopods (*Neogonodactylus oerstedii* n=25) were purchased from a private distributor and collected along the Florida Keys. Marine crustaceans were shipped directly to the Arizona laboratory aquaria where they were held in marine tanks maintained with filtered water that approximated ocean salinity (approx. 35 ppt). Freshwater crayfish were kept in tanks with filtered RO (reverse osmosis) water. Aquaria were kept at room temperature and maintained on a 12-hour light/dark cycle. Animals were fed a diet of bloodworms and dried shrimp pellets.

Additional specimens were collected and maintained at Friday Harbor Laboratories (University of Washington; Friday Harbor, WA). Spiny lebbeids (*Lebbeus groenlandicus* n=37), squat lobsters (*Munida quadrispina* n=2), northern crangon (*Crangon franciscorum* n=10), *Ligia pallasii*, *Nebalia pugettensis*, Dana’s blade shrimp (*Spirontocaris lamellicornis* n=3), and horned shrimp (*Paracrangon echinata* n=8) were collected by trawling along the Puget Sound near San Juan Island. Ghost shrimps (*Neotrypaea californiensis* n=11), northern hooded shrimp (*Betaeus harrimani* n=3), and purple shore crabs (*Hemigrapsus nudus* n=40) were collected along shorelines near Friday Harbor, as were hairy hermit crabs (*Pagurus hirsutiusculus* n=35). All animals collected at Friday Harbor were kept alive in shaded outdoor tanks with constantly circulating seawater until time of processing.

### Antibodies

A variety of antibodies were used in this study to visualize neural architecture and identify regions of neural connectivity involved with learning and memory. Table 1 contains a comprehensive list of all antibodies used here, as well as their source, target, concentration, and supplier.

**Table 1.**
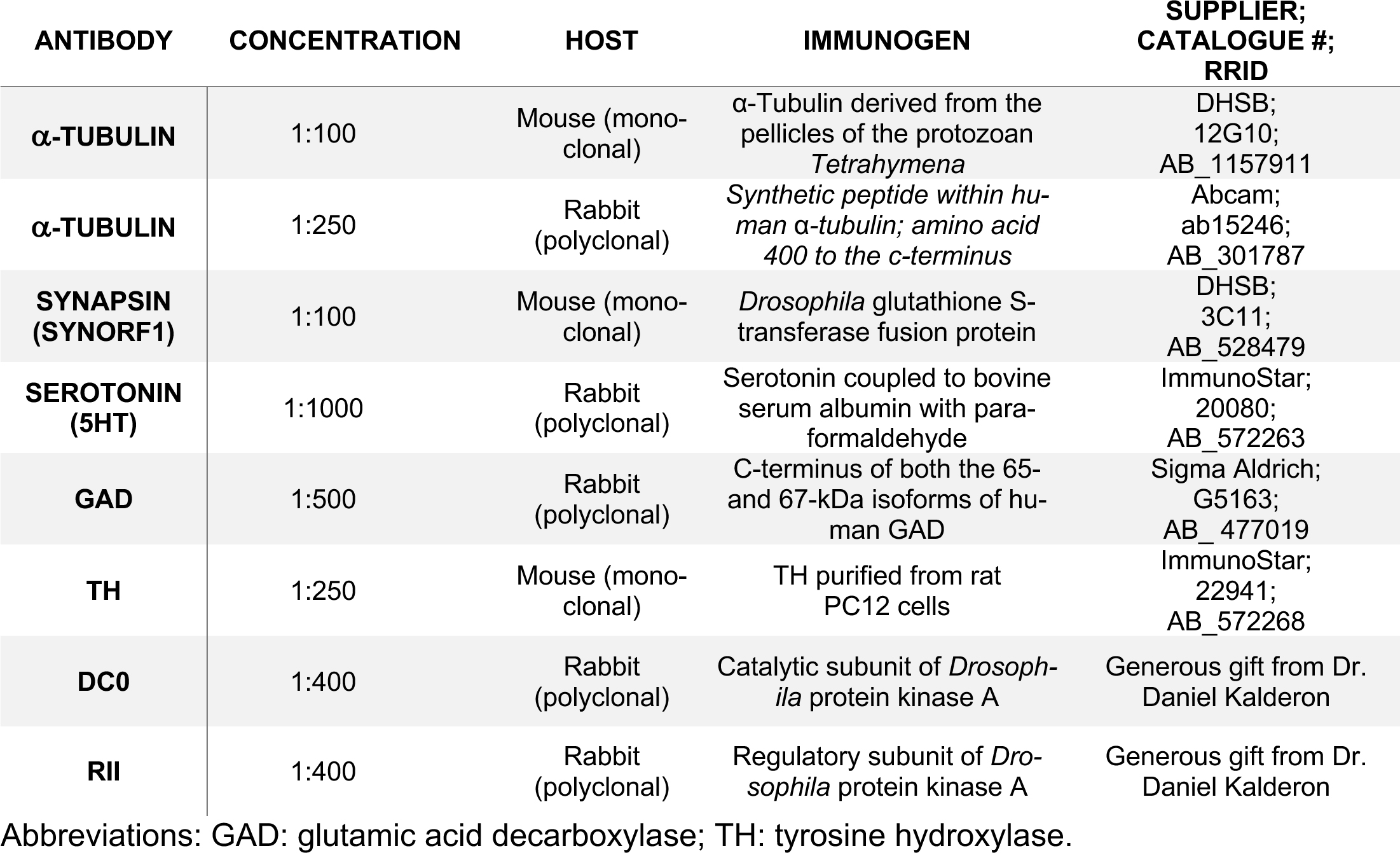
Antibodies used for this study.

To detect regions of synaptic densities and cytoskeletal elements, antibodies against synapsin and *α*-tubulin were used, respectively. Immunostaining against synapsin was carried out using a monoclonal antibody raised against *Drosophila* GST-synapsin fusion protein (SYNORF1; Klagges et al., 1996). Two different antibodies were used to label the cytoskeletal protein, *α*-tubulin. The first was raised in mice against the protozoan *Tetrahymena α*-tubulin (Thazhath et al., 2002). The second was generated in rabbit against human *α*-tubulin, and was used to allow co-labelling in experiments in which the other primary antibody was raised in mouse. Antibodies against synapsin and *α*-tubulin have been widely used as neural markers across distant phyla, suggesting that both proteins have been highly conserved across evolutionary time (Harzsch et al., 1997; Harzsch and Hansson, 2008; Stemme et al., 2012; Sullivan et al., 2007; Brauchle et al., 2009; Andrew et al., 2012). Additional antibodies raised against serotonin (5HT), glutamic acid decarboxylase (GAD), and tyrosine hydroxylase (TH) were used to further aid in the visualization of neural arrangements. Serotonin is a ubiquitous neurotransmitter, and has previously been used across invertebrate phyla as a marker for phylogenetic analyses (Harzsch and Waloszek 2000; Harzsch, 2004; Stemme et al., 2012; Dacks et al., 2006). GAD and TH are enzymatic precursors to gamma-aminobutyric acid (GABA) and dopamine, respectively, and were used in this study to detect putative GABAergic and dopaminergic neurons. Anti-GAD has previously been used to identify GABAergic neurons in remipedes (Stemme et al., 2016) and crustaceans (Wolff et al., 2017; Sayre and Strausfeld, 2019). Similarly, anti-TH has been used as a marker to detect putative dopaminergic neurons. Co-labelling experiments have revealed that anti-GAD and anti-TH reliably label GABAergic or dopaminergic cells when used in conjunction with primary antibodies against either GABA or dopamine, respectively (Cournil et al., 1994; Crisp et al., 2001; Stern, 2009; Stemme et al., 2016).

Antibodies against DC0 and RII were used to detect regions likely to be involved with learning and memory. These antibodies recognize the major regulatory (RII; Li et al., 1999) and catalytic (DC0) subunits of the *Drosophila* c-AMP-dependent enzyme, protein kinase A (PKA; Lane and Kalderon 1993; Skoulakis et al., 1993; Crittenden et al., 1998). Anti-RII and anti-DC0 label identically, and Western blot analysis has shown that anti-DC0 reliably labels the expected band across remipedes, crustaceans, and insects (Wolff et al., 2012; Stemme et al., 2016).

### Immunohistochemistry

Animals were anesthetized by cooling on ice. Heads bearing both eyestalks were removed and placed immediately in freshly made ice-cold fixative containing 4% paraformaldehyde (PFA; Electron Microscopy Sciences; Cat# 15710; Hatfield, PA) and 10% sucrose in 0.1 M phosphate-buffered saline (PBS; Sigma Aldrich; Cat# P4417; St. Louis, MO). Eyestalk and midbrain neural tissue were dissected free and allowed to fix overnight at 4°C.

The following day, tissue was rinsed twice in PBS and transferred to Eppendorf tubes containing a preheated agarose-gelatin solution consisting of 7% gelatin, 5% agarose. The tissue was left in this mixture and kept warm in a 60°C water bath for 1 hour (adapted from Long, 2018). Tissue was then transferred to a plastic mold containing fresh embedding media and allowed to solidify at 4°C for 5-10 min. Next, blocks were removed from the mold, trimmed, and post-fixed in 4% PFA in PBS for 1 hour at 4°C. Blocks were then rinsed twice in PBS and vibratome sectioned at 60 μm (Leica; Nussloch, Germany). Following sectioning, tissue slices were rinsed twice in PBS containing 0.5% Triton-X (PBST; Electron Microscopy Sciences; Cat# 22140; Hatfield, PA) and blocked in a solution of 5% normal donkey serum (Jackson ImmunoResearch; RRID:AB_2337258; West Grove, PA) for 1 hour. The primary antibody was then added and the sections were left overnight on a gentle shaker at room temperature (RT). TH immunolabelling required an alternative immunostaining method, as is described below. Control experiments in which the primary antibody was not added resulted in the complete absence of immunostaining.

The next day, sections were rinsed 6 times over the course of an hour in PBST. Meanwhile, 2.5 μl of secondary antibodies were added to Eppendorf tubes at a concentration of 1:400 and centrifuged at 11,000 *g* for 15 minutes. Secondary IgG antibodies used in this study were raised in donkey and conjugated to either Cy3, Cy5, or Alexa 647 fluorophores (Jackson ImmunoResearch; RRID: AB_2340813; RRID: AB_2340607; RRID: AB_2492288, respectively; West Grove, PA). After centrifugation, the top 900 μl from the secondary antibody solution was added to the tissue sections, and the sections were left to soak on a gentle shaker overnight at RT. Following secondary antibody treatment, sections were rinsed twice in 0.1M Tris-HCl (Sigma Aldrich; Cat# T1503; St. Louis, MO) buffer. To label cell nuclei, SYTO™ 13 (Thermo Fisher Scientific; Cat# S7575; Waltham, MA) were used at a concentration of 1:2000 and left to incubate for 1 hour on a vigorous shaker. Lastly, tissue sections were washed in Tris-HCl for 6 changes over the course of an hour before being mounted in a medium consisting of 25% glycerol, 25% polyvinyl alcohol, and 50% PBS using #1.5 coverslips (Fisher Scientific; Cat# 12-544E; Hampton, NH).

In some preparations phalloidin conjugated to Alexa 488 (Thermo Fisher Scientific; RRID: AB_2315147; Waltham, MA) was used to label the cytoskeletal protein, f-actin. Sections processed this way were incubated in a solution containing the conjugated phalloidin at a concentration of 1:40 in PBST for 2-3 days following secondary antibody treatment. Sections were left in the dark on a gentle shake at RT for the duration of this time, and then subsequently mounted as described above.

For TH immunolabelling, neural tissue was treated using a much shorter fixation time of just 30-45 minutes. Longer fixation periods resulted in a reduction or complete loss of immunostaining, as has been previously reported in locusts (Lange and Chan, 2008). Additionally, antibody labelling was best preserved when carried out on whole mounts before sectioning (previously reported in Cournil et al., 1994). Following fixation, whole mount tissue was rinsed twice in PBS and then twice in PBST over the course of 20 minutes. Brains were then blocked as described above for 3 hours on a gentle shake. Primary antibody solution was added to the tissue, and the preparations were exposed to microwave treatment. This included two cycles of 2 minutes low power, 2 minutes no power, under a vacuum at 18°C. Whole brains were then left for 2-3 days in primary antibody solution on a gentle shake at RT and were subjected to microwave treatment each day of antibody incubation. Following primary antibody treatment, tissue was rinsed 6 times in PBST for 1 hour and left to incubate for 1-2 days in secondary antibody (secondary antibodies were centrifuged as described above). Afterwards, the whole mounts were rinsed 6 times over the course of an hour, embedded, sectioned, and mounted as described above.

### Bodian silver staining

Nerve fibers were resolved using Bodian’s reduced silver stain (Bodian, 1936). Nervous tissue was dissected in acetic alcohol formalin fixative (AAF; 10 ml 16% paraformaldehyde, 2.5 ml glacial acetic acid, 42 ml 80% ethanol) and left to fix for 3 hours at RT. Tissue was rinsed twice in 50% ethanol and then transferred through an ethanol series of 70%, 90%, and two changes of 100% ethanol for 10 minutes each. Next, dehydrated tissue was soaked in *α*-terpineol (Sigma Aldrich; Cat# 432628; St. Louis, MO) for 15 minutes before being swapped into xylenes for an additional 15 minutes. Tissue was then embedded in paraffin, sectioned at 12 μm, and processed following Bodian’s original method (Bodian, 1936; for further details see Sayre and Strausfeld, 2019). It is important to note that one of the most deterministic staining variables in this method is the silver proteinate, more commonly known by its commercial name, Protargol (Bodian, 1936; Pan et al., 2013). To the dismay of many anatomical biologists, effective forms of Protargol have been commercially discontinued in the past decade. Other forms of silver proteinate that have become commercially available since the disappearance of Protargol have proven to be ineffective for this staining procedure (Pan et al., 2013). For the present study, silver proteinate was generated in-house, following the method described in Pan et al., 2013 (Figures 4C, 5A, D-E, 6F, 7C). The exceptions are Figure 2D and 9A which used the now unobtainable French “Roche Proteinate d’Argent”, and Figure 10A which was processed using Protargol; Winthrop Chemical Co., NY).

### Golgi impregnations

Animals were anesthetized over ice, and midbrain and eyestalk neural tissue was dissected free from the exoskeleton and enveloping sheath in an ice-cold fixative solution containing 1 part 25% glutaraldehyde (Electron Microscopy Sciences; Cat# 16220; Hatfield, PA) to 5 parts 2.5% potassium dichromate (Sigma Aldrich; Cat# 207802; St. Louis, MO) with 3-12% sucrose in high pressure liquid chromatography (HPLC)-grade distilled water (Sigma Aldrich; Ca#270733). Dissected whole brains were then transferred to fresh fixative and left overnight in a dark cupboard at RT. The rest of the protocol was also carried out at RT, with preparations kept in the dark whenever possible. The use of metal tools was avoided at all stages following the dissection to prevent contamination with the highly reactive silver nitrate.

The next morning, brains were briefly rinsed in 2.5% potassium dichromate and placed in a solution of 2.5% potassium dichromate and 0.01% osmium tetroxide (Electron Microscopy Sciences; Cat# 19150; Hatfield, PA) for around 12 hours. After osmification, neural tissue was post-fixed for 24 hours using freshly made 1:5 admixture of 25% glutaraldehyde:2.5% potassium dichromate, omitting sucrose.

Tissue was then washed in 2.5% potassium dichromate three times for 10-minute intervals before silver impregnation: brains were quickly transferred three times from the potassium dichromate solution to glass containers containing fresh 0.75% silver nitrate (Electron Microscopy Sciences; Cat# 21050; Hatfield, PA) in HPLC water and then left overnight in silver nitrate. The next day, brains were rinsed twice in HPLC water. Some tissue was dehydrated and embedded as described below. For mass impregnation of neurons, brains were treated with a double impregnation in which the osmification step followed by a second treatment of silver nitrate was repeated.

After silver impregnation, brains were rinsed twice in HPLC water and dehydrated through a four step ethanol series from 50% to 100% ethanol at 10-minute intervals. Ethanol was then replaced by propylene oxide (Electron Microscopy Sciences; Cat# 20401; Hatfield, PA), and tissue was left for 15 min before being transferred through an increasing concentration of Durcupan embedding medium (Sigma Aldrich; Cat# 44610; St. Louis, MO) in 25%, 50%, and 75% propylene oxide over the course of 3 hours. Tissue was left overnight in 100% liquid Durcupan and then polymerized in BEEM capsules at 60°C for 12-24 hours. Cooled blocks were sectioned at a thickness of 20-40 μm and mounted using Permount mounting medium (Fisher Scientific; Cat# SP15-100; Hampton, NH).

## RESULTS

In the following, descriptions of mushroom bodies and their evolved derivatives are organized as sections arranged to match the oldest to most recent lineages shown in Figure 1A,B. Figure 1B provides a summary of anti-DC0immunoreactive territories described as homologies of the mushroom body ground pattern. Each description is preceded by a brief commentary on the species considered.

**Figure 1.**
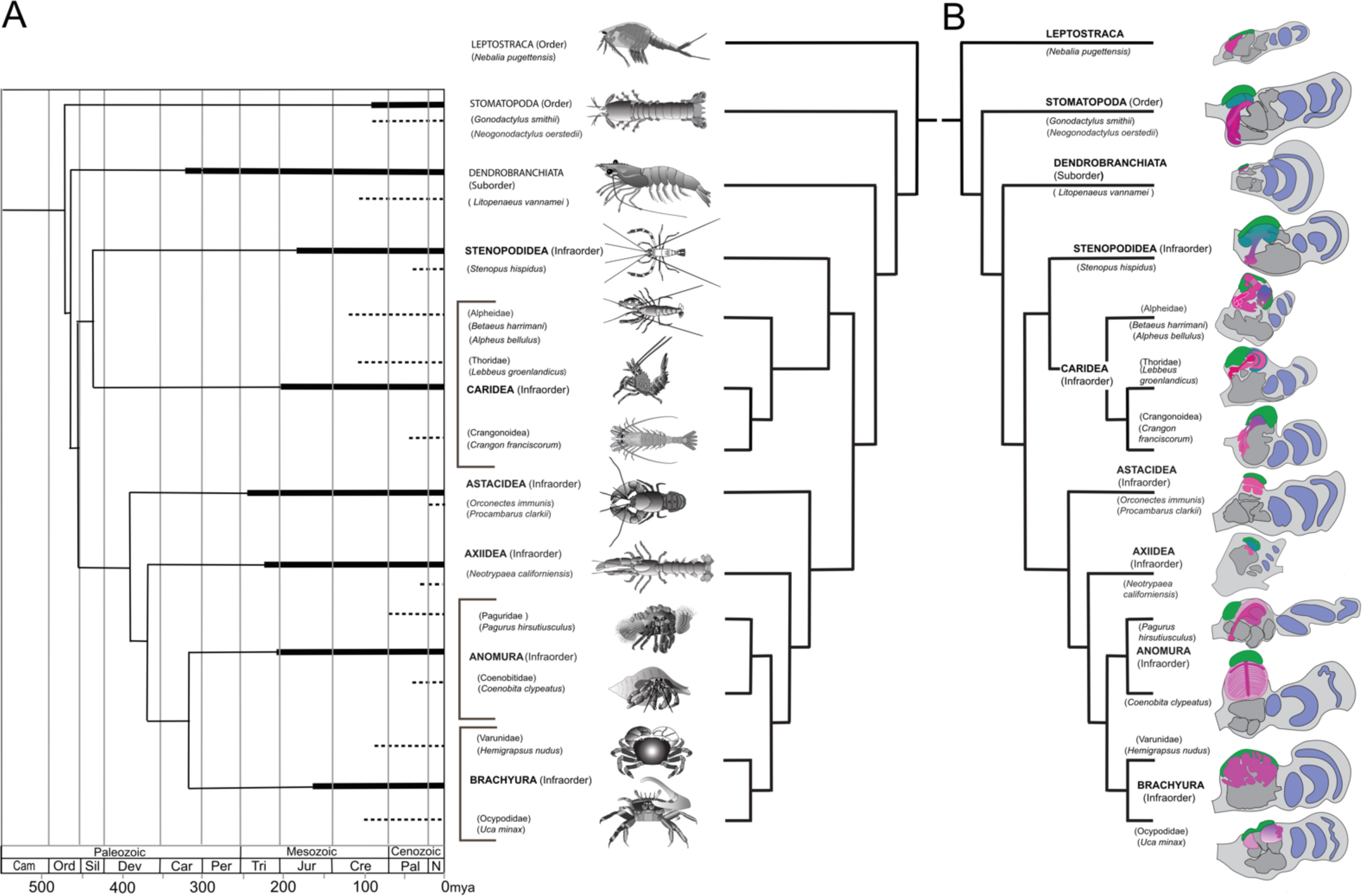
Time lines, phylogeny and protocerebral morphology of malacostracan lineage representatives. Time lines and lineage relationships are based on the molecular phylogeny of Wolfe et al. (2019). Geological time scale shown as millions of years ago (mya). Images depict species used for this study, including *Nebalia pugettensis* representing the phyllocarid sister of Eumalacostraca. **A**. Solid lines indicate estimated occurrence of lineages sampled for this study; dashed lines indicate estimated age of representative taxon (see citations in text). **B**. Schematic showing actual proportions of anti-DC0-immunoreactive centers (shades of magenta) in the right lateral protocerebrum of species described in this account. Rostral is up, distal to the right. Nested optic lobe neuropils shown light blue; rostrally disposed globuli cell clusters, green; generalized neuropil domains of the lateral protocerebrum, mid-gray.

### The evolutionary timeline

Species considered here belong to eumalacostracan lineages whose divergence times are known from fossil-calibrated molecular data (Wolfe et al., 2019), and which are estimated to have originated some time between the mid-to-late Ordovician and the Carboniferous (Figure 1A). The following descriptions include only a small number of the species belonging to lineages used for molecular phylogenomic reconstruction (see Schwentner et al., 2018; Wolfe et al., 2019). We have also referred to analyses that focus on mantis shrimps (Van der Wal et al., 2017), as well as decapods, including stenopods (cleaner shrimps), alpheids and carideans (“visored shrimps”, “pistol shrimps”: Anker et al., 2006; Anker and Baeza, 2012; Bracken et al., 2010; Davis et al., 2018), brachyurans (“true crabs”: Tsang et al., 2014), thalassinids (“ghost shrimps”: Tsang et al., 2008), anomurans including “hermit crabs” (Bracken-Grissom, 2014a; Chablais et al., 2011), and various clades that colloquially are referred to as lobsters (see Shen et al., 2013; Bracken-Grissom, 2014b). Figure 1B summarizes the disposition and shapes of anti-DC0immunoreactive centers in their lateral protocerebra.

The principal organization of the olfactory pathway of crown Crustacea up to the level of the lateral protocerebrum is reminiscent of that in Hexapoda. However, claims of homology (Schachtner et al., 2005) are insecure (see Discussion and Table 2). With the exception of Cephalocarida (Stegner and Richter, 2011), axons of relay neurons from olfactory centers in the deutocerebrum ascend rostrally as two prominent mirror-symmetric fascicles, called the olfactory globular tracts (OGT). These axonal pathways are massive, often comprising many thousands of axons. Left and right OGTs converge in the mid-protocerebrum, just dorsal to the central body where most if not all their axons bifurcate sending a tributary into both lateral protocerebra (Figure 2). In Eumalacostraca, the lateral protocerebral neuropils are usually resident within the enlarged volume of the eyestalks’ terminal articles. Exceptions are in land hermit crabs, where the lateral protocerebra reside at the base of the eyestalks. In lineages that lack eyestalks (species of Alpheidae, pistol shrimps, hooded shrimps) and Thalassinidae (ghost shrimps) or lack compound eyes (e.g., Copepoda and Remipedia), the lateral protocerebra immediately flank the protocerebrum’s midbrain.

**Figure 2.**
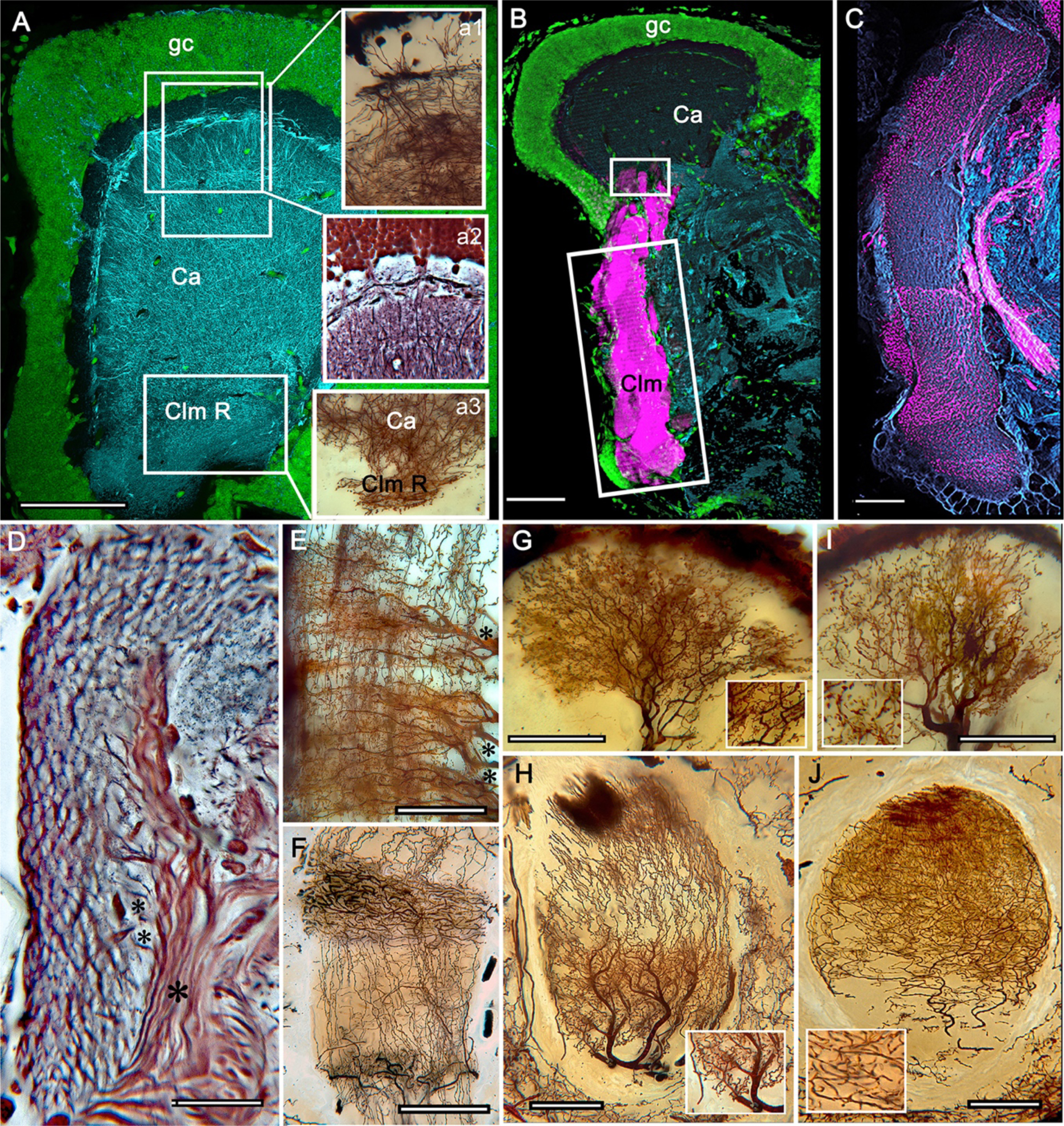
The stomatopod mushroom body: *Neogonodactylus oerstedii*. **A.** Immunohistochemical labelling of basophilic globuli cells (gc, green, syto13) that give rise to neurites (cyan, anti-*α*-tubulin), the dendritic trees of which form the calyx (Ca). **a1** Golgi impregnation reveals individual globuli cells, with their neurites projecting into the dense calycal meshwork below; **a2** Bodian staining shows stratifications of intrinsic neuron dendrites at the outer boundary of the calyx; **a3** Golgi impregnation of axon-like processes of intrinsic neurons at their convergence at the base of the Ca where they form the root of the columnar lobe (Clm R). **B.** Confocal laser scan showing strong labelling of anti-DC0 (magenta) along the length of the largest of the mushroom body’s columnar lobes (Clm; cyan, anti-*α*-tubulin; green, syto13). The small rectangle denotes the origin of the column from the calyx, as shown in panel A, inset a3. The larger rectangle corresponds to views of the same column using different techniques, shown in panels C and D. **C.** Antityrosine hydroxylase (magenta) and anti-*α*-tubulin (cyan) immunolabelling reveals discrete synaptic domains of mushroom body output neurons (MBON) along the length of a column. **D.** Bodian-stained section corresponding to long rectangular inset of panel B. Parallel fibers reminiscent of insect mushroom body (MB) lobes interweave along the length of a MB column. Bold asterisk marks a bundle of neurons whose processes contribute to synaptic domains through the length of the column. Small asterisks mark individual processes as they separate out from the much larger bundles. **E,F.** Golgi-impregnated afferent and efferent processes (asterisks in E) innervate discrete domains along the stomatopod (E) MB column and, for comparison the column of an insect MB (F, cockroach). **G,H.** Cross sections of MB columns in a stomatopod (G) and cockroach (H) demonstrate spine-like specializations (insets) of dendrites that typify MBONs. **I,J.** Cross sections of terminal arborizations invading the lobes of a stomatopod (I) and cockroach (J) MB; insets show corresponding varicose and beaded presynaptic specializations. Scale bars in A, 50μm; B, 100μm; C-G, 50μm; H, 100μm; I, 50μm; J, 100μm.

**Table 2.** Correspondence of main morphological characters defining the crustacean and insect olfactory pathways. Characters 1-7 pertain to the deutocerebrum. Characters 10-12 pertain to the protocerebrum.

| Character | Hexapoda (Dicondylia) | Malacostraca and Remipedia | Correspondence |
| --- | --- | --- | --- |
| **1. Ligand-gated odorant receptors (ORs)** | Ubiquitous | None | – |
| **2. Ionotropic olfactory receptor channels (IRs)** | Present | Ubiquitous | + |
| **3. Olfactory neurons in each sensillum** | 1-5 (in Dicondylia) | Numerous, >10* and can exceed 100s | – |
| **4. Olfactory receptor neuron axon** | Targets one genetically corresponding glomerulus | Not targeting: can innervate multiple glomeruli | – |
| **5. Olfactory lobe subunits** | Unique glomeruli relating to genetic receptor identity | Discrete subunits, isomorphic: number unrelated to receptor identities | – |
| **6. Accessory lobes** | None | Reptantain lineages (possible remipede homologue) | – |
| **7. OL Projection neurons (PNs)** | Most uniglomerular | All multiglomerular | – |
| **8. Accessory lobe Projection neurons (via OGT)** | Not applicable | In Reptantia | – |
| **9. Number of PN axons from olfactory lobe** | 2–5 X number of glomeruli | Some thousands, unrelated to glomerulus number | – |
| **10. Projection neuron axon tract (OGT or AGT)** | AGT ipsilateral | OGT bifurcates, ipsi- and contralateral. | – |
| **11. Protocerebral PN terminals** | In MB calyx + other lateral protocerebral targets | In MB calyx + other lateral protocerebral targets | + |

### Ground pattern and derived organization of the protocerebral olfactory center

Uniquely identifiable neuropils in the lateral protocerebrum include a prominent anti-DC0immunoreactive center, the intrinsic neurons of which originate exclusively from rostrally located clusters of minute basophilic neuronal perikarya. In Decapoda, such anti-DC0-immunoreactive centers can adopt three morphologies: a columnar neuropil subtending a robust cap of synaptic neuropil (also referred to as a calyx), together providing a mushroom body (Figure 2); a prominent neuropil protruding rostrally that lacks a lobe and may be divided into two or more distinct territories, referred to as a hemiellipsoid body (Figure 2); or a broad folded domain extending within almost the entire rostral half of the lateral protocerebrum. This latter arrangement may be specific to Brachyura (crabs; see Figure 1B). Extremely small anti-DC0-immunoreactive centers may represent evolved reductions (Figure 3). All of these centers receive as their major afferent supply olfactory relay neurons from the brain’s second segment, the deutocerebrum. As in insects, in stomatopods mushroom bodies have been reported as integrating multimodal sensory information (Thoen et al., 2019) as have hemiellipsoid bodies in achelate reptantians (Mellon, 2000; Mellon et al., 1992; McKinzie et al., 2003).

**Figure 3.**
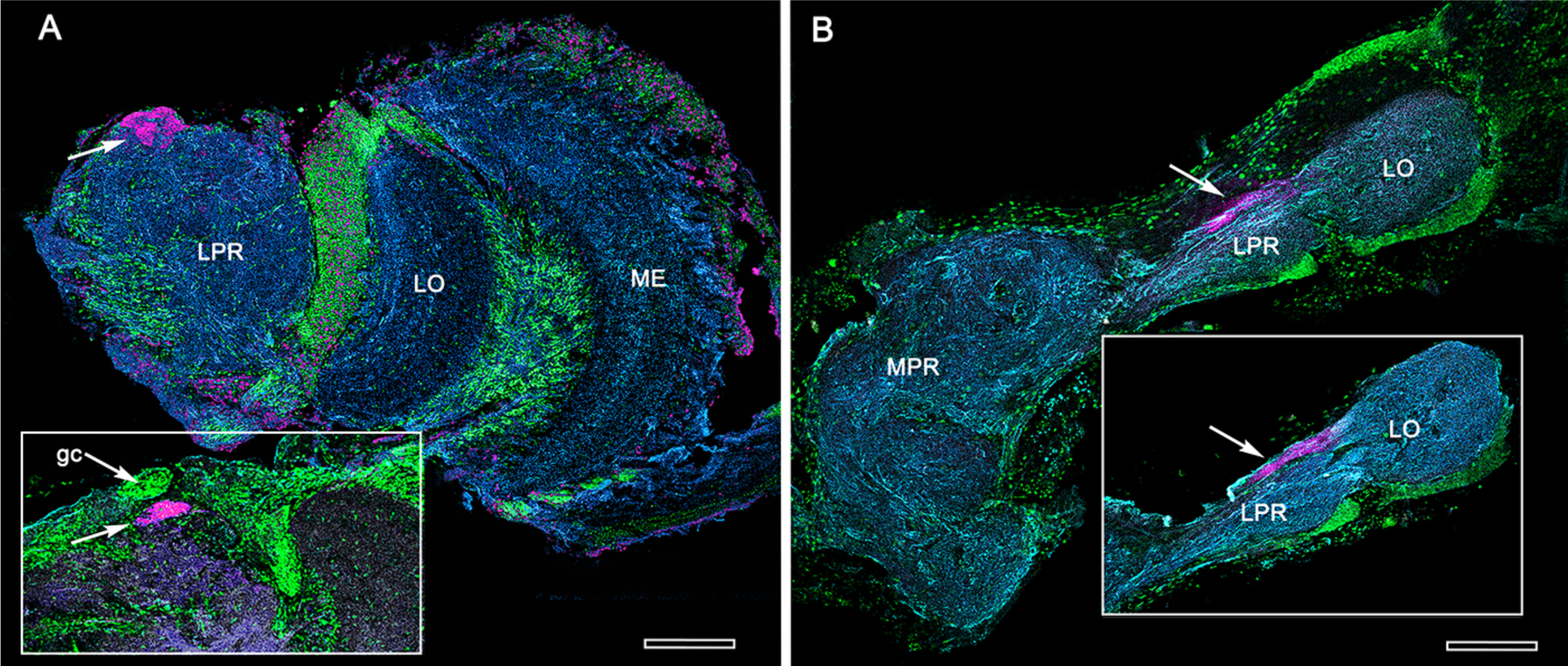
Extreme reduction of DC0-immunoreactive neuropils in decapods and isopods. **A**. The pelagic white leg shrimp *Penaeus vannamei*, showing substantial lateral protocerebral neuropils (LPR), large optic neuropils (ME, medulla; LO, lobula), but a minute anti-DC0-immunoreactive center the location of which corresponds to that of the ancestral mushroom body of Stomatopoda and Caridea, and the derived hemiellipsoid body of Reptantia. Inset: Another example showing also the clustered globuli cells (gc) overlying the immunoreactive center. **B**. Preparations from two individual isopods *Ligia pallasii*, showing the medial protocerebrum (MPR) connected to a greatly reduced lateral protocerebrum (LPR). A narrow layer of anti-DC0-immunoreactive neuropil resides at a position corresponding to that occupied by the mushroom body of a stomatopod or the hemiellipsoid body of a reptantian. Scale bars,100 μm.

### The stomatopod mushroom body

To introduce the crustacean mushroom body we briefly recap and expand our findings from Stomatopoda (Figure 2; see also, Wolff et al., 2017). Stomatopoda is sister to all other Eumalacostraca. Its lineage has been claimed as extending back to the Devonian, represented by fossil Tyrannophontidae (Hof, 1998), but the relationships of this fossil to extant stomatopod morphology is highly problematic. Unambivalent fossil Stomatopoda are Cretaceous (Wolfe et al, 2016) and their age by molecular clock data is Triassic (Van der Wal et al., 2017).

In Stomatopoda, the mushroom body’s intrinsic neurons originate from adjoining clusters of basophilic cell bodies (globuli cells). Two clusters contribute to a calyx comprising two contiguous territories, from which arise two parallel columnar extensions. Together, with two other columns originating from two additional clusters of globuli cells associated with reduced or absent calyces, these impart a quadripartite organization corresponding to that of an insect mushroom body (Wolff et al., 2017). Further correspondences are the arrangements of intrinsic neurons in the stomatopod calyces, the organization of calycal microglomeruli, their axon-like extensions of intrinsic cells in the lobes, and the division of the lobes into discrete synaptic domains. Notably, intrinsic neuron dendrites define at least three discrete layers through the calyces (inset a1, a2, Figure 2A). Their axon-like prolongations converge at the base of the calyces to form the columnar lobes (inset a3, Figure 2A). In Stomatopoda and in all dicondylic insects, the lobes are intensely labelled by antibodies raised against DC0 (Figure 2B), the *Drosophila* orthologue of vertebrate PKA-C*α*, which in vertebrates and invertebrates plays a crucial role in synaptic facilitation (Burrell and Sahley, 2001; Abel and Nguyen, 2008). Numerous other characters defining the stomatopod mushroom body correspond to those resolved in a basal dicondylic insect, exemplified by the cockroach *Periplaneta americana,* as well as in more recently diverged groups such as Drosophilidae. Synaptic domains partitioning the columnar lobes are defined by modulatory and peptidergic efferent and afferent processes (Figure 2C). The processes of intrinsic neurons form characteristic orthogonal “Hebbian” networks (Figure 2D), and these orthogonal networks are further defined by afferent and efferent neurons that intersect them (Figure 2E). The cross-sectional profile of the column may vary across species – it is oval in *Periplaneta* for example, but more discoid in the stomatopod – the spinous and varicose attributes of afferent and efferent neurons are identical (Figure 2G-J).

### Reduced protocerebral centers in Dendrobranchiata

Dendrobranchiata is the second oldest Eumalacostracan lineage today represented by pelagic penaeid shrimps. Using antibodies raised against D-tubulin, FMRFamide, and allatostatin on *Penaeus vannamei*, Meth et al. (2017) were unable to delineate a target neuropil supplied by the olfactory globular tract (OGT). Their assumption was, however, that there had to be a hemiellipsoid body. Sullivan and Beltz (2004), using dye tracing into the OGT of *Penaeus duorarum,* had identified a small twinned neuropil situated extremely rostral in the lateral protocerebrum. The neuropil receives collaterals from the OGT, the axons of which mainly terminate in lateral protocerebral neuropil beneath. Observations of *Penaeus vannamei* (Figure 3A), also a peripatetic/pelagic dendrobranchiate, identifies an anti-DC0immunoreactive neuropil, corresponding to the location identified by Sullivan and Beltz (2004). Immunohistology resolves a small intensely immunoreactive four-component center. A similarly reduced center typifies some isopods (Figure 3B). Later in the Discussion we suggest that these findings demonstrate evolved reduction and loss of these centers in pelagic free-swimming taxa or ones that live in ecologies with a restricted range of odorants.

### Stenopodidid mushroom bodies

Mitochondrial genomics resolve the infraorder Stenopodidea as the sister taxon of Caridea (Shen et al., 2013). Its molecular clock age is Jurassic with a Cretaceous fossil calibration (Wolfe et al., 2019). The following description is restricted to the coral banded cleaner shrimp *Stenopus hispidus*, one of the few well-documented cleaner shrimps that inhabit tropical coral reefs and groom fish of external parasites or detritus (Vaughan et al., 2017). These shrimps are well known for their knowledge of place, occupying a specific territory – their cleaning station – to which they return daily and which are visited by favored fish (Limbaugh et al. 1961). Long-term memory in *S. hispidus* is suggested by observation of individuals pair bonding and recognizing a partner even after absences of several days (Johnson, 1977). Allocentric recognition of place and partner recognition are mediated by chemoreception (Esaka et al., 2016).

We confirm here Sullivan and Beltz’s (2004) original description of the stenopid *S. hispidus* lateral protocerebral center that they referred to as a hemiellipsoid body, divided into three domains that receive layered OGT terminals. However, this center should not be designated as a hemiellipsoid body because reduced silver stains demonstrate that the three domains (Ca1-Ca3) assume the configuration of a mushroom body calyx (Figures 4, 5), with each domain contributing many hundreds of parallel axon-like fibers from intrinsic neurons (Figure 5A,D,G) that converge into a common root from which arises a compound columnar lobe terminating as three grouped tubercular domains (Figures 4A-C,E; 5A,B). Parallel fibers branch in these volumes where they provide preand postsynaptic specializations (Figure 4A,B). Immunolabelling with anti-DC0 demonstrates this protein expressed at very high levels in the tubercles and at lower levels just distally in the columnar lobe (Figure 4D-F). And, as shown by immunolabelling with antibodies against GAD, 5HT and TH, the tubercles are further divided into smaller territories such that the entire structure assumes the serial domain arrangement characterizing the mushroom body ground pattern (Figure 4G-J).

**Figure 4.**
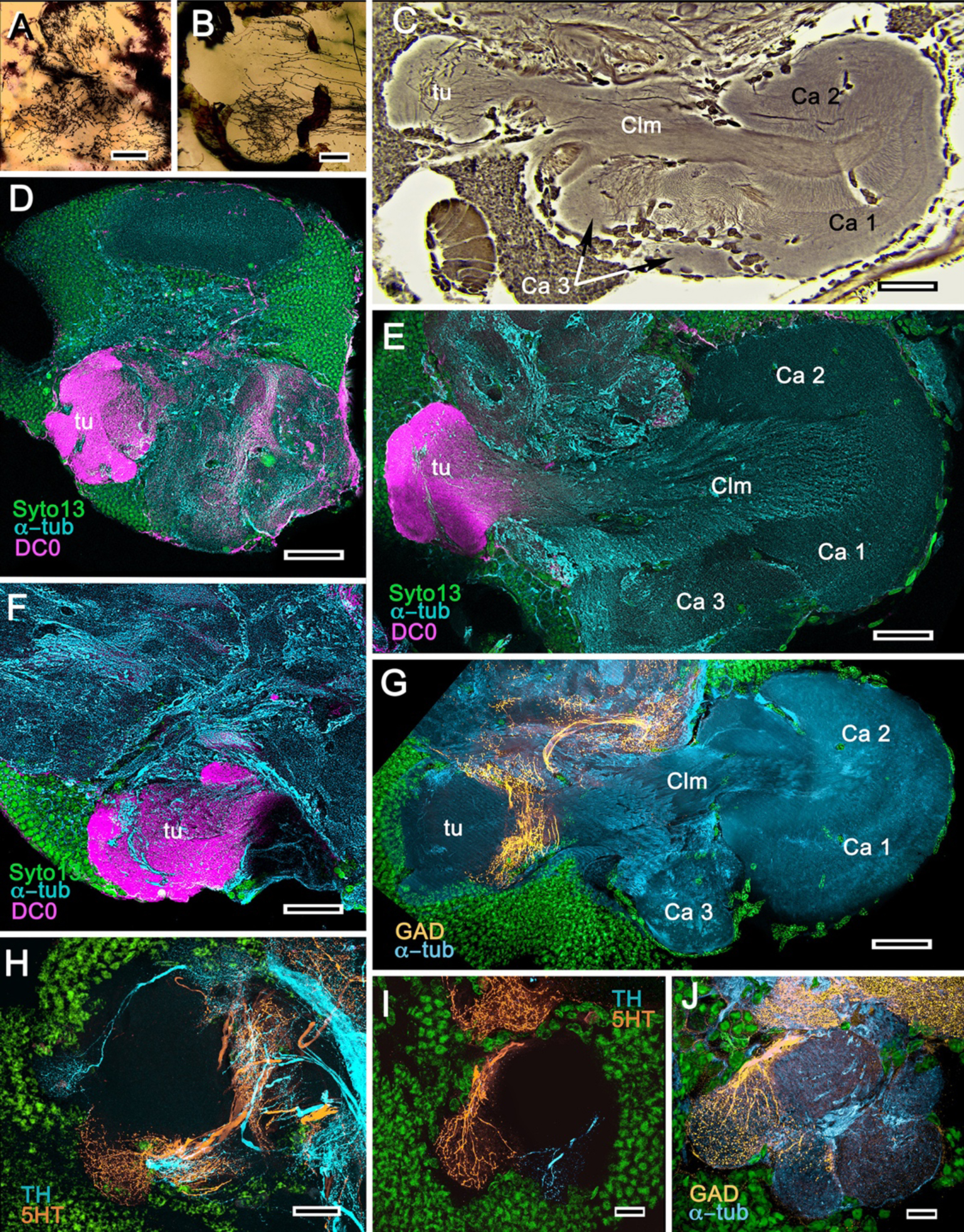
Mushroom bodies of *Stenopus hispidus*: DC0 and neuromodulatory delineation of tubercular domains. **A,B.** Golgi-impregnated sections of neurites arborizing in the mushroom body column tubercle. **C.** Bodian silver stained overview of the stenopid mushroom body. Axon-like extensions from three distinct calycal regions (Ca1, Ca2, Ca3) provide the column (Clm) that terminates in discrete tubercular domains (tu). **D-F.** Immunolabelled sections showing regions containing elevated levels of anti-DC0 (magenta). Anti-DC0 immunoreactivity dominates the entirety of the column’s tubercular domain (cyan, *α*-tubulin; green, syto13). **G.** Calycal domains and their columnar extension show no anti-GAD immunoreactivity (yellow), with the exception of a few processes that innervate specific zones within the tubercle (green, syto13). **H,I.** Anti-5HT (serotonin, orange) and anti-tyrosine hydroxylase (TH, cyan) immunoreactive fibers similarly occupy discrete territories in the column’s tubercle (green, syto13). **J.** Tubercular domains showing discrete territories, thus subsets of intrinsic neuron projections, occupied by anti-GAD-immunoreactive processes (yellow; cyan, *α*-tubulin; green, syto13). Scale bars in A, B, 20μm; C, 50μm; D-H, 50μm; I, J, 20μm.

The calyces, which alone correspond to Sullivan and Beltz’s “hemiellipsoid body”, are composed of many thousands of intrinsic neuron dendrites. Their axon-like extensions, resolved by Golgi impregnations (Figure 5A), converge predominantly beneath Ca2 in the columnar root but also recruit from both Ca1 and Ca3 through an elaborate system of interweaving processes (upper and lower insets, Figure 5A). Groups of axons segregate before invading different tubercles, each denoting a discrete integrative domain.

**Figure 5.**
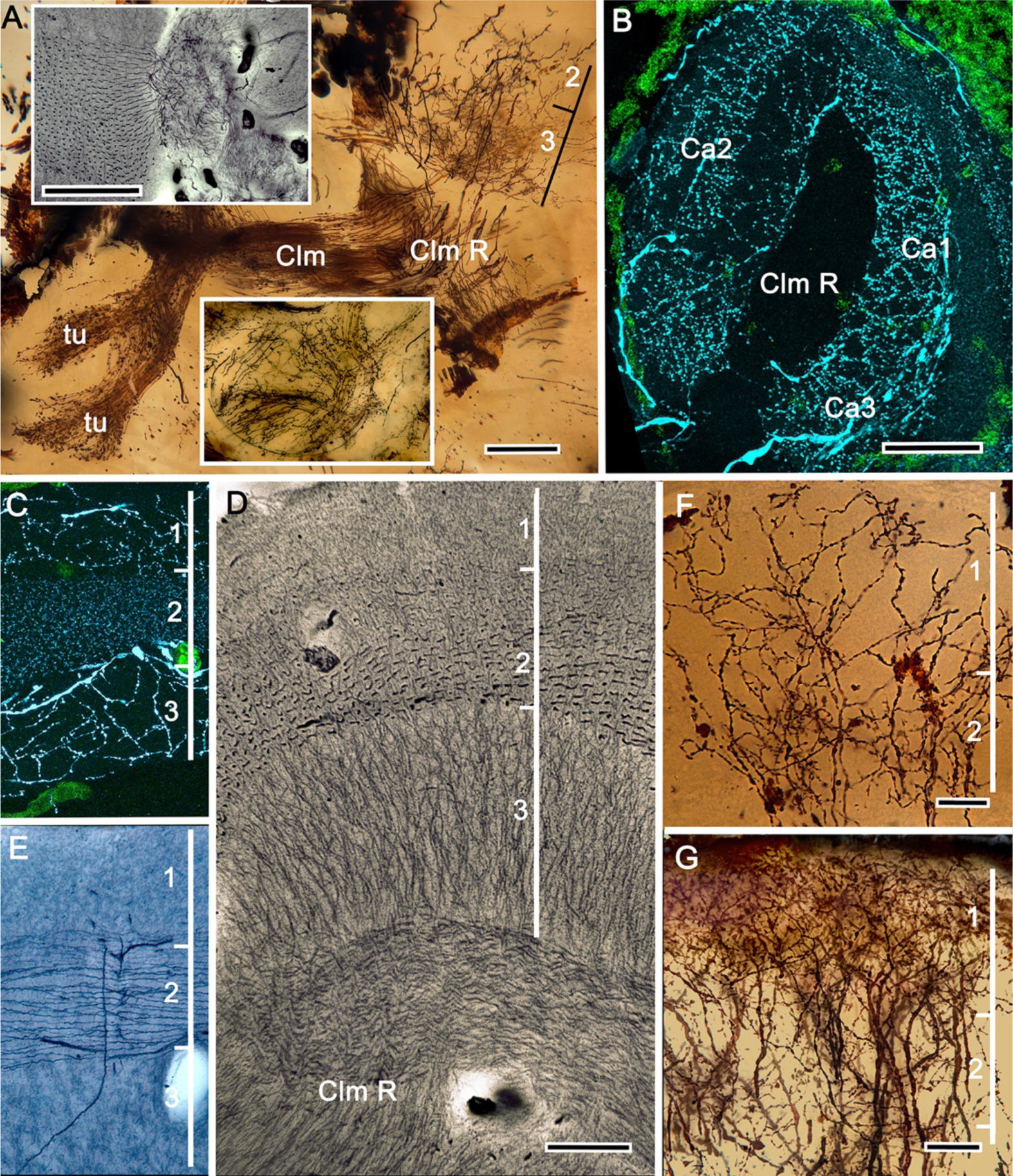
The stenopid mushroom body: cytoarchitecture of the calyx. **A.** Mushroom body intrinsic neurons extend their axon-like processes from the calycal domain (upper inset) into the column (Clm) terminating in its tubercles (tu, bottom inset). 2, 3 indicate the two inner layers of the calyx. **B.** Anti-tyrosine hydroxylase immunolabelling (TH, cyan; cell bodies, green) reveals the supply by anti-TH-immunoreactive processes to all three calyces surrounding the root of the mushroom body column (Clm R). **C.** Enlargement of the calyx showing anti-TH-positive neuronal arborizations in calycal layers 1 and 3, while anti-TH labelling in layer 2 shows very fine, yet densely populated neuron varicosities. **D.** Bodian stain showing the stratified stenopid calyx. Layer 1 contains the apical dendrites of mushroom body intrinsic neurons, as well the terminals of afferent neurons from other lateral protocerebral centers. Terminals of the olfactory globular tract (OGT) arborize in layer 2. Layer 3 is occupied by the extensions of the inner dendrites of intrinsic neurons. Intrinsic neurons send their collaterals into the root of the columnar lobe (Clm R; also see **A**, top inset) after extending laterally through layer 3. **E.** Bodian stain of layer 2, which is further defined by numerous parallel projecting terminals from the olfactory lobes carried by the OGT. **F,G.** Golgi impregnations of slender, varicose afferent neurons (**F**) and apical dendrites of intrinsic neurons (**G**) in layer 1 of the stenopid calyx. Scale bars In A, 20μm; B, 50μm; C-E, 20μm; F, 100μm; G, 50μm.

As in *Lebbeus* and Stomatopoda, the calyces are profusely invaded by branches of TH-containing neurons (Figure 5B). These are arranged through three successive calycal layers (Figure 5C-D), each of which is defined by its elaborate cytoarchitecture reflecting specific neuronal components, as described from Stomatopoda and the alpheoid shrimp *Lebbeus* (Wolff et al., 2017; Sayre and Strausfeld, 2019). Layer 1 of each of the three calyces (Ca1-Ca3) contains ascending processes of slender varicose afferents and the apical dendrites of intrinsic neurons (Figure 5G, F). Climbing fibers enter the calyces from other lateral protocerebral centers. Layer 2 (Figure 5D, E) is denoted by the numerous parallel projecting terminals originating from the olfactory globular tract. Layer 3 (Figure 5D) contains the massively arborizing inner dendrites of intrinsic neurons, some of which are shown at the corresponding level in Figure 5A, upper right. Processes of intrinsic neurons sweep laterally beneath layer 3 to enter the root of the mushroom body’s columnar lobe (Clm R; Figure 5D).

### Caridean mushroom bodies

Caridea spilt from their sister group Procaridea in the Carboniferous (Bracken et al., 2010), with most caridean crown group diversification occurring in the Jurassic (Wolfe et al., 2019). Caridea comprises the second largest decapod taxon, with two-thirds of its species occupying marine habitats. Here, we consider marine species belonging to the families Alpheidae and Thoridae (members of the superfamily Alpheoidea), and intertidal species of the family Crangonidae. These families originated approximately 220, 160 and 70 mya, respectively (Davis et al., 2018). Alpheidae is a diverse family that contains snapping shrimps, commonly referred to as pistol shrimps, which include the only known crustaceans to have evolved eusociality (Duffy et al., 2000), and visored shrimps closely related to the genus *Alpheus* (Anker et al., 2006). Pistol shrimps are represented here by the non-social species *Alpheus bellulus*, which, like other snapping shrimps, possesses asymmetric claws (chelae). The visored shrimp is *Betaeus harrimani*. Like pistol shrimps, *B. harrimani* is an active predator, although not employing concussion, and is also facultatively commensal, sharing burrows of the ghost shrimp *Neotrypaea californiensis* (Anker and Baeza, 2012). Unlike the genus *Alpheus*, *Betaeus* has symmetric claws, reflecting the more basal position of this genus within Alpheidae (Anker et al., 2006). Comparisons of the brains of *A. bellulus* and *B. harrimani* revealed almost no discernible differences other than that the eyes and visual neuropils of *B. harrimani* are smaller than those of *A. bellulus*. Figures 6 and 7 therefore illustrate relevant features of the mushroom bodies using both species.

**Figure 6.**
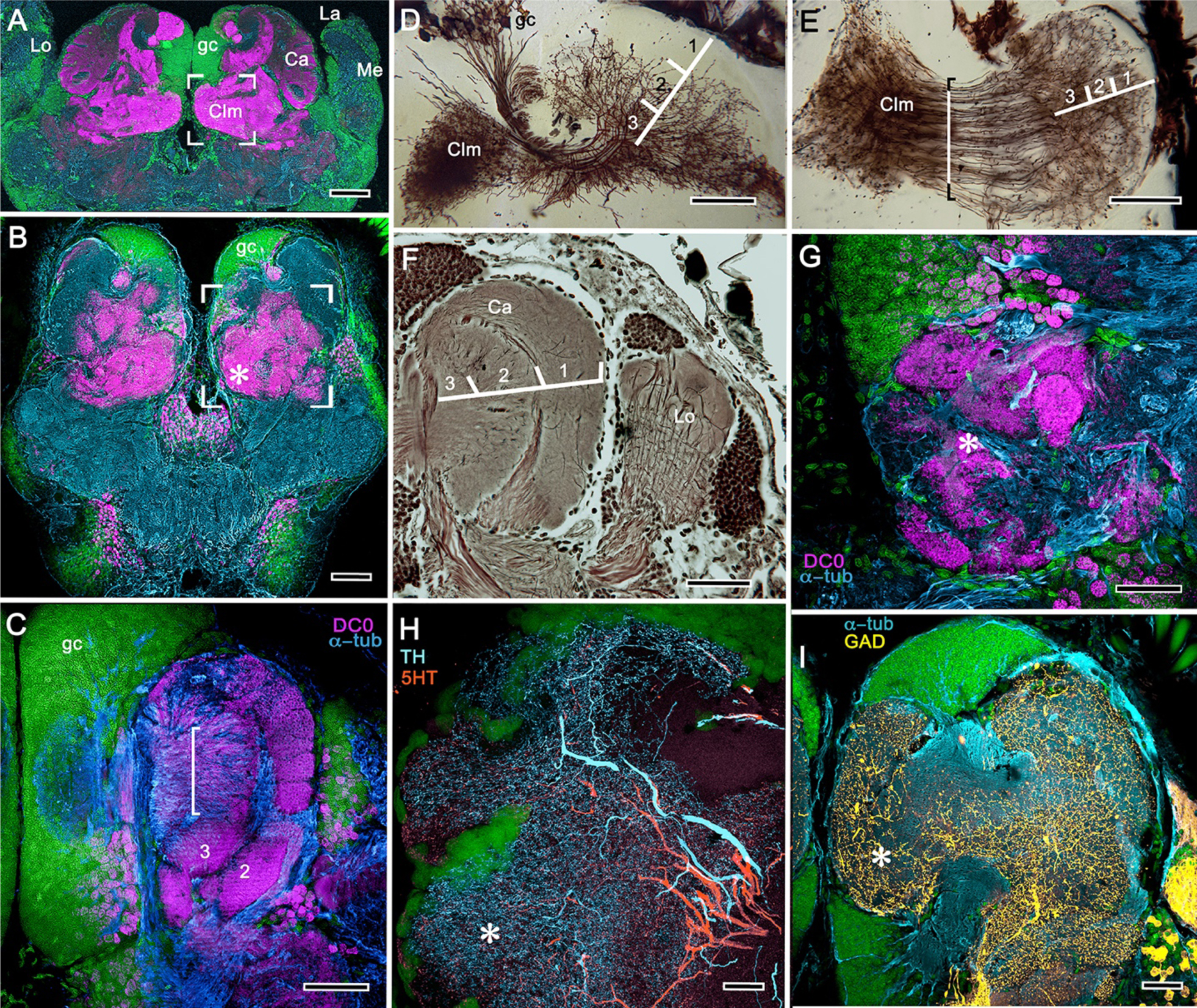
Neuroarchitecture of the alpheid mushroom body. **A.** Anti-DC0 (magenta) immunoreactivity in the brain of *Alpheus bellulus*. The mushroom bodies (Ca calyx; Clm columnar lobe) are flanked on both sides of the brain by the optic lobes (cyan, anti-*α*-tubulin; green, syto13; La, lamina; Me, medulla; Lo, lobula). **B.** Anti-DC0 (magenta) immunolabelling in brain of *Betaeus harrimani.* Asterisk marks columnar lobe ending in tubercles (cyan, anti-*α*-tubulin; green, syto13). **C.** Anti-DC0 expression in the mushroom body calyx of *A. bellulus*. Throughout its depth, and in each layer (layers 2 and 3 are shown here), the calyx shows a high affinity for antibodies raised against DC0. Striations (bracket) correspond to bundles of intrinsic neuron processes supplying the columnar lobe, correspondingly bracketed in panel E. **D,E.** Golgi impregnations of mushroom body intrinsic cell clusters, originating from globuli cells (gc) and giving rise to dendrites in the calyces (Ca, layers numbered 1-3) and column (Clm). The distinct calycal layers are most clearly revealed in Bodian-stained sections (**F**). **G.** Anti-DC0 is expressed in distinct territories within the *A. bellulus* tubercle. **H,I.** Aminergic processes in the lateral protocerebrum of *B. harrimani*. Asterisks in G, H, I denote corresponding regions. **H.** Anti-5HT (orange) and anti-TH (cyan) show these neural arborizations invading most regions of the lateral protocerebrum coincident with volumes denoted by anti-DC0-labelled mushroom body-associated structures. **I.** Anti-GAD (yellow) immunoreactivity shows an expression pattern throughout the anti-DC0-positive domains (cyan, anti-*α*-tubulin). Scale bars in A (*A. bellulus*), 100μm; B (*B. harrimani*), 100μm; C-F (*A. bellulus*), 100μm; G (*A. bellulus*), 40μm; H (*B. harrimani*), 20μm; I (*B. harrimani*) 50μm.

Immunolabelling with anti-DC0 reveals an elaborate system of neuropils expressing high levels of the antigen, demonstrating the relationship of those areas predominantly to the globuli cell cluster medially and to the protocerebrum rostrally (Figure 6A, B). Notably, alpheid species lack eyestalks, and possess very small compound eyes. The nested optic centers (lamina, medulla and lobula) are correspondingly reduced, and the enormous mushroom bodies occupy much of the rostral and lateral protocerebrum’s volume (Figure 6A, B). The calyx is elaborate, showing through its depth high affinity to DC0 (Figure 6C). As in Stomatopoda and Stenopodidea, the calyx has three layers, each of which is defined by the dendritic disposition of its intrinsic cells (Figure 6D-F). Intrinsic cells extend their axon-like processes in parallel bundles for a short distance before forming elaborate networks in the columnar lobes (Figure 6D, E), which extend dorsoventrally as convoluted volumes comprising tubercular domains (Figure 6G). These volumes are heavily invested by anti-TH-, anti-5HT- and anti-GAD-positive arborizations (Figure 6H-I).

As in Stomatopoda and varunid Brachyura (shore crabs), the alpheid protocerebrum has a prominent DC0-positive reniform body, the components of which correspond to those described for *Neogonodactylus oerstedii* and *Hemigrapsus nudus* (Thoen et al., 2019). This center consists of four discrete neuropils linked by a branching axonal tract (pedestal) that extends axons from a glomerular initial zone to three separate zones (lateral, distal and proximal) each with its own characteristic immunomorphology (Figure 7A-E). Abutting the mushroom body ventrally, these territories are similar to, but spatially distinct from a columnar extension from the calyx that, like the main columnar ensemble shown in Figure 6B,G, consists of discrete tubercles, each denoted by its affinity to antisera against GAD, DC0, TH and 5HT (Figure 7F-I).

**Figure 7.**
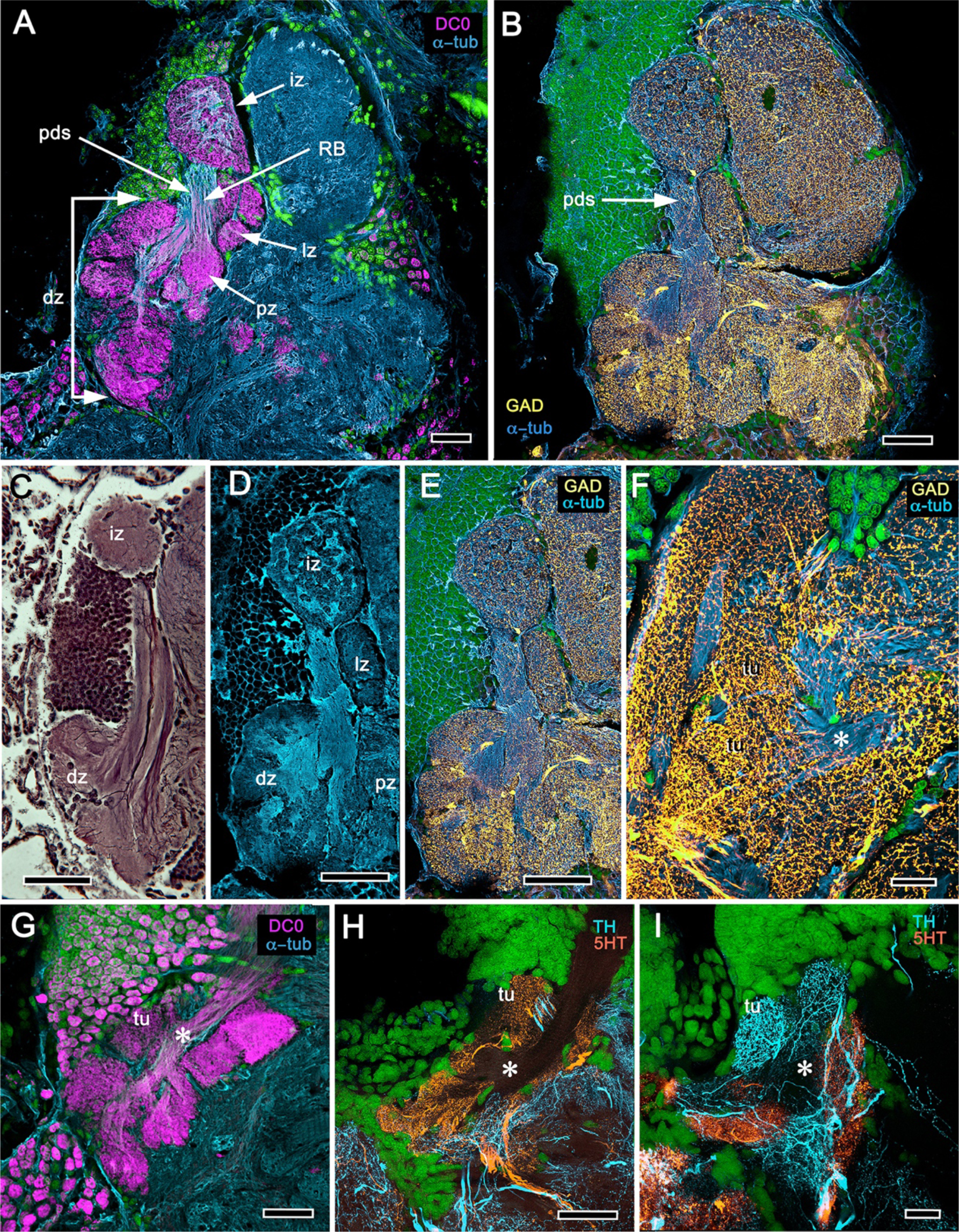
Reniform body (A-E) and mushroom body lobes (F-I) in alpheid shrimps. **A-E.** Neuroarchitecture of the *Alpheus bellulus* reniform body (RB; cyan, anti-*α*-tubulin; green, syto13). **A.** Components of the alpheid reniform body are clearly distinguished by their high affinity for anti-DC0 (magenta). A pedestal-like (pds) bundle of neurites give rise to four distinct zones: the initial zone (iz), the proximal zone (pz), the distal zone (dz) and the lateral zone (lz). **B,E.** Anti-GAD immunolabelling (yellow) reveals putative inhibitory processes occupying all major zones of the reniform body. **C,D.** Bodian silver-stained *Betaeus harrimani* (**C**) and anti-*α*-tubulin-labelled *A. bellulus* sections (**D**) reveal the nearly identical compartmental layout of the reniform body across the two alpheid species. **F-I.** Anti-DC0 and aminergic innervation of discrete domains within the mushroom body (MB) columnar lobe. Asterisks in panels F–I indicate the MB column just before its processes defasciculate into discrete tubercular domains. These are shown in panels G-H where one corresponding tubercle (tu) is indicated. **F.** Innervation of tubercular domains by anti-GAD-immunoreactive processes (yellow). **G.** Anti-DC0 labelling is shown expressed within tubercles. **H,I.** Glomerular divisions in the tubercular domain are delineated by anti-TH- and anti-5HT-immunoreactive fibers. In all panels green indicates syto13-stained neuronal cell bodies. Scale bars in A-C,E, 50μm (*B. harrimani*); D,G-I, 50μm; F, 10μm (*A. bellulus*).

We next consider Thoridae, a separate lineage of the superfamily Alpheoidea that includes the genera *Lebbeus* and *Spirontocaris*.

*Lebbeus groenlandicus* and *Spirontocaris lamellicornis* are exotically patterned and protected from predation by sharp spiny protuberances from their exoskeleton. They are solitary, highly active, and territorial, living on reefs or rocks. The two species look similar; both are patterned into irregular white and orange-pink bands, a feature suggesting self-advertisement or aposomatic warning. There is one report of a lebbeid cleaning the mouths of fish, reminiscent of behavior by the banded coral shrimp described above (Jensen, 2006). The present description of their mushroom bodies (Figures 8,9,10A) expands a recent study of *Lebbeus* (Sayre and Strausfeld, 2019), resolving two completely distinct calyces that share a common organization into three concentric layers. These correspond to the dense arrangements of intrinsic neurons (inset to Figure 8A). Defined by their immunoreactivity to DC0 and affinity to a palette of antibodies (Figure 8A,B,D), the layers in the larger calyx (Ca1, Figure 8A) demonstrate elaborate synaptic strata enriched with synaptic proteins (Figure 8C). Processes resolved by anti-5HT correspond to intrinsic neuron layers, as do dense arrangements of processes belonging to 5HT-positive afferent neurons (Figure 8B, C) and arrangements of anti-TH-positive terminals (Figure 8D). Thin processes of intrinsic neurons belonging to the two calyces extend centrally as two separate columnar lobes.

**Figure 8.**
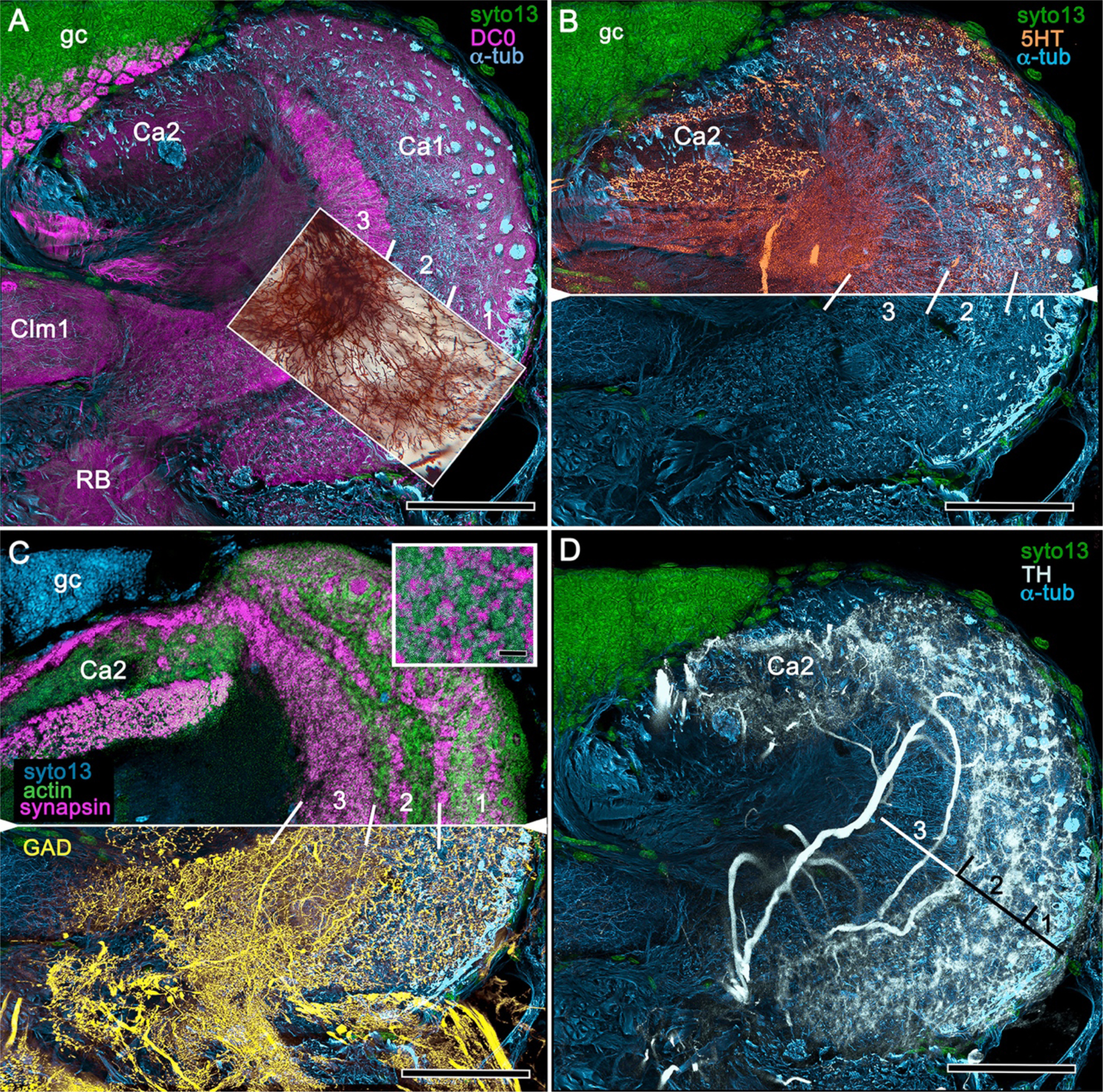
Neuroanatomy of mushroom body calyces in the thorid shrimp, *Lebbeus groenlandicus*. **A-D**. Confocal laser scans of immunohistochemically labeled sections (anti-*α*-tubulin is cyan in all panels except C). **A.** Anti-DC0 (magenta) immunoreactivity in both calycal regions (Ca1, calyx 1; Ca2, calyx 2), as well as in the columnar extension of intrinsic cells (Clm1) and the reniform body (RB). The inset demonstrates the palisade of Golgi-impregnated intrinsic cells, the dendritic organization of which reflects the presence of three distinct layers in Ca1 (1-3 against inset). **B.** Anti-5HT immunolabelling showing afferent neuron terminals from various regions in the lateral protocerebrum ending in both calyces, and the three layers in Ca2 (upper half of panel). Calycal cytoarchitecture is distinguishable by anti-*α*-tubulin labelling alone (lower half of panel). **C.** Upper half. Double labelling with anti-synapsin and f-actin further resolves layering (1-3) of synaptic sites in both Ca1 and Ca2. Inset. High-resolution scans resolve synaptic microglomeruli. Lower half. Anti-GAD-immunopositive fibers (yellow) extending from the lateral protocerebrum to provide dense innervation into all calycal levels. **D.** Large-diameter anti-TH-positive terminals (white) spread their tributaries throughout calycal layers 1 and 2. Scale bars in A-D, 100μm.

Clm1, the column provided by Ca1, demonstrates an orthogonal network – characteristic of insect and stomatopod mushroom bodies – of intrinsic cell processes intersected by afferent and efferent neurons and by modulatory elements (Figure 9B-E,H,I). The columns contain dense arrangements of synaptic specializations (Figure 9F,G) and although these do not clearly distinguish successive synaptic territories, the arrangement of anti-TH-positive processes reveal this organization, one of the diagnostic characteristics of the mushroom body ground pattern (Figure 9I). The smaller of the two calyces provides intrinsic fibers to its columnar extension (Clm2), which terminates as a system of tubercles corresponding to the organization of the mushroom bodies in basal insects (Zygentoma, Ephemeroptera). These also show dense arrangements of synaptic sites (Figure 9J, inset).

**Figure 9.**
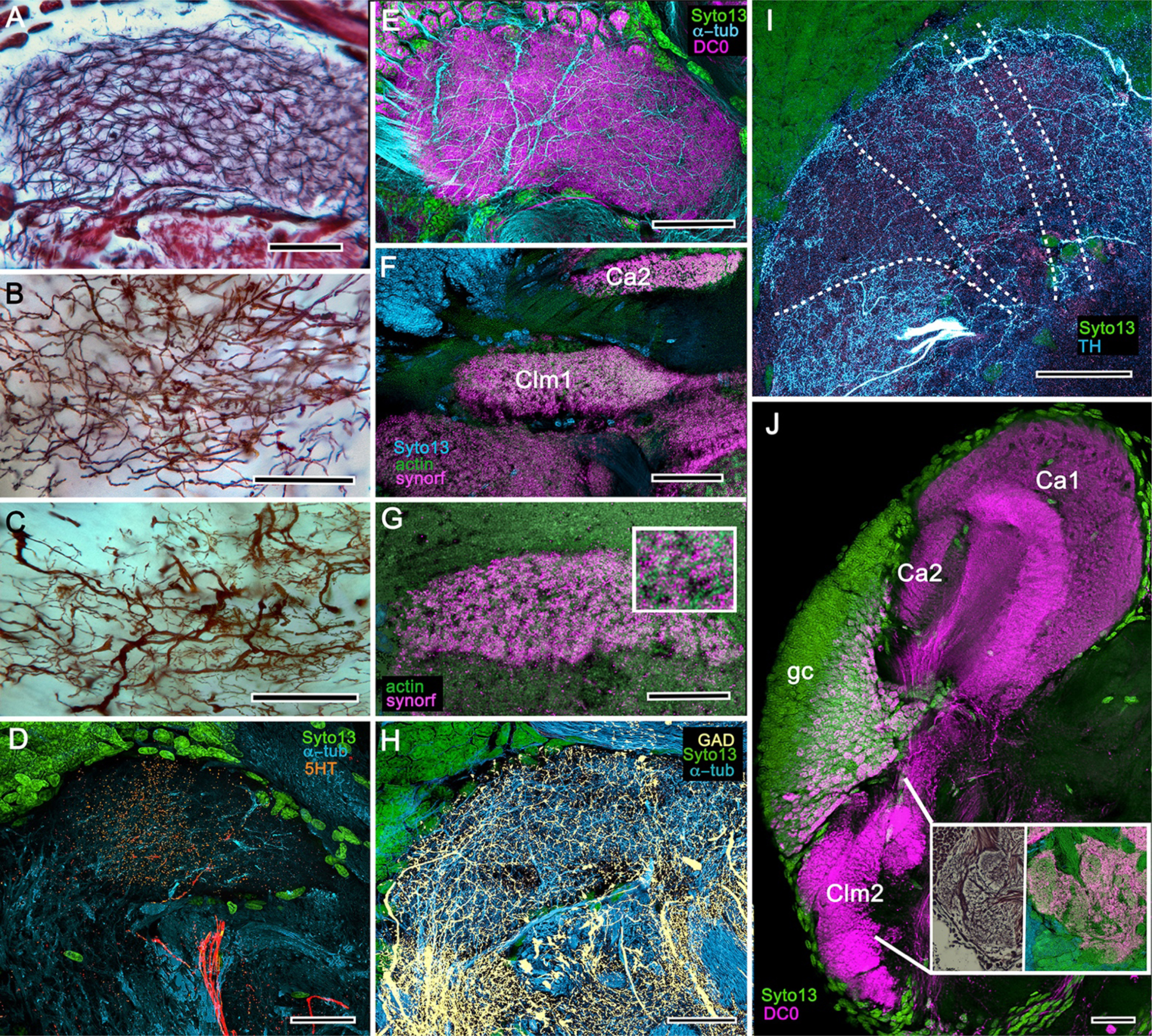
Columnar projections of *Lebbeus* mushroom body intrinsic neurons. **A.** Bodian-stained extensions of intrinsic neurons interweave throughout the length of the first column (Clm1). **B**. Golgi impregnations resolve these decorated by bouton- and spine-like specializations denoting, respectively, afferent terminals and mushroom body output neurons (MBONs). **C.** Golgi impregnations resolve afferent and efferent (MBON) processes intersecting serial domains within Clm1. **D.** Sparse anti-5HT-immunoreactive neurons send terminals into Clm1 from regions in the lateral protocerebrum. **E.** Anti-DC0 (magenta) immunoreactivity reveals a high affinity for this antibody in Clm1 where it outlines afferent and *α*-tubulin-positive efferent fibers. **F,G.** Anti-synapsin (magenta) expression in Clm1 and Ca2. Double labelling with anti-synapsin (SYNORF1) and f-actin reveals microglomeruli in Clm1 (**G**, inset). **H.** Anti-GAD-positive processes project along the full length of Clm1. **I.** Discrete serial domains along Clm1 are defined by processes of MBONs immunoreactive to anti-TH (cyan). **J.** Anti-DC0 immunoreactivity defines Clm2 showing its characteristic tubercles (compare with Fig. 10A). The two insets resolve tubercles in Bodian silver-stained material (left) and anti-synapsin/f-actin labelling of their synaptic zones (right). Scale bars in A-J, 50μm.

The genus *Crangon* is here represented by the sand shrimp *Crangon franciscorum* and the horned shrimp *Paracrangon echinata*, belonging to the superfamily Crangonoidea, which originated at the end of the Cretaceous (Davis et al., 2018). Both species provide interesting comparisons with the alpheids with respect to their habitat and mode of life. *Paracrangon* looks superficially like an almost colorless version of *Lebbeus*. However, unlike *Lebbeus,* which is highly active and hunts on rocky substrates and reefs, *P. echinata* is an almost immobile nocturnal ambush predator confined to crevasses. Its relative inactivity may be attributable to having only four walking legs compared to *Lebbeus*‘s six (Jensen, 2011). The lateral protocerebrum of *P. echinata* (Figure 10C) is radically different from that of *S. lamellicornis* (Figure 10B) or *Lebbeus* (Figure 10A). It possesses a single but substantial calyx that has an intense affinity to anti-DC0, the cytoarchitecture of which appears to be generally homogeneous apart from a subtle indication of a peripheral layer (Figure 10C). It gives rise to a diminutive system of anti-DC0-positive parallel fibers that provide a narrow column terminating in a small tubercular cluster (Figure 10D). Of all the caridean species investigated, the modified mushroom body of *P. echinata* most suggests convergence with the reptantian achelate-type morphology (see Discussion).

**Figure 10.**
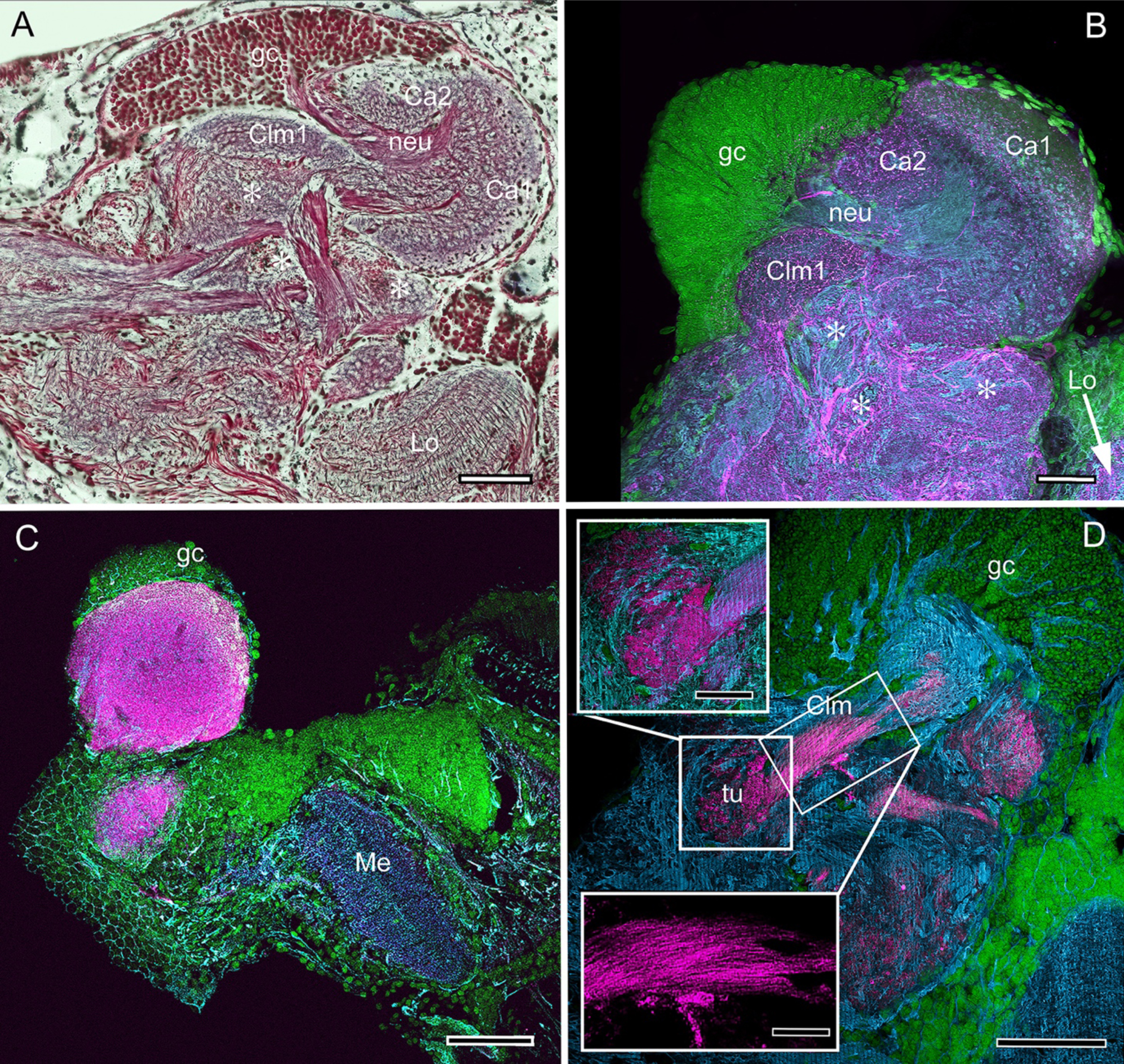
Mushroom body neuroarchitecture: *Spirontocaris lamellicornis* and *Paracrangon echinata*. **A.** Bodian-stained section of *Lebbeus groenlandicus* demonstrates that the organization of the mushroom body calyces and columns of this thorid shrimp is almost identical to that of species sharing the same ecological niche, exemplified by the thorid *S. lamellicornis* and the crangonidid *Paracrangon echinata*. **B**. Anti-*α*-tubulin (cyan) and anti-GAD immunoreactivity (magenta) in *S. lamellicornis* resolve the two calyces Ca1, Ca2, and their neurites (neu) leading from globuli cells (gc; green, syto13) and Clm1. More caudal regions of the corresponding lateral protocerebrum are indicated by asterisks in panels A and B. **C.** Anti-DC0 immunostaining of *P. echinata* reveals high levels of the antigen in a substantial rounded calyx with a narrowly differentiated outer stratum covered by a dense layer of globuli cells (gc). **D.** A second specimen showing evidence of a diminutive column (Clm), resolved by anti-DC0, comprising parallel fibers (inset, lower left) ending in a small tubercular (tu) domain (inset, upper left). Me medulla; Lo lobula. Scale bars in A-D, 100μm; insets in D, 50μm.

The last caridean considered here is *Crangon franciscorum,* which inhabits subtidal sandy habitats where it actively preys on smaller shrimp species, amphipods and other microfauna (Siegfried, 1982). The calyx of *C. franciscorum* shows low affinity to anti-DC0 but gives rise to a prominent system of anti-DC0positive columns that terminate in swollen tubercles (Figure 11A). The entire system of the calyx, lobes and adjacent neuropils, including the visual system’s lobula, is densely packed with anti-GAD-positive processes (Figure 11B). The calyx possesses stratified arborizations, resolved by anti-5HT and anti-TH immunostaining, some of which suggest subdivisions laterally across the calyx (Figure 11C,D). In this species, several of the lateral protocerebral neuropils caudal to (beneath) the calyx appear to comprise many neuropil islets, each with as strong an affinity to anti-DC0 as has the calyx. The disposition of these islets suggests a system possibly corresponding to the reniform body (Figure 11A).

**Figure 11.**
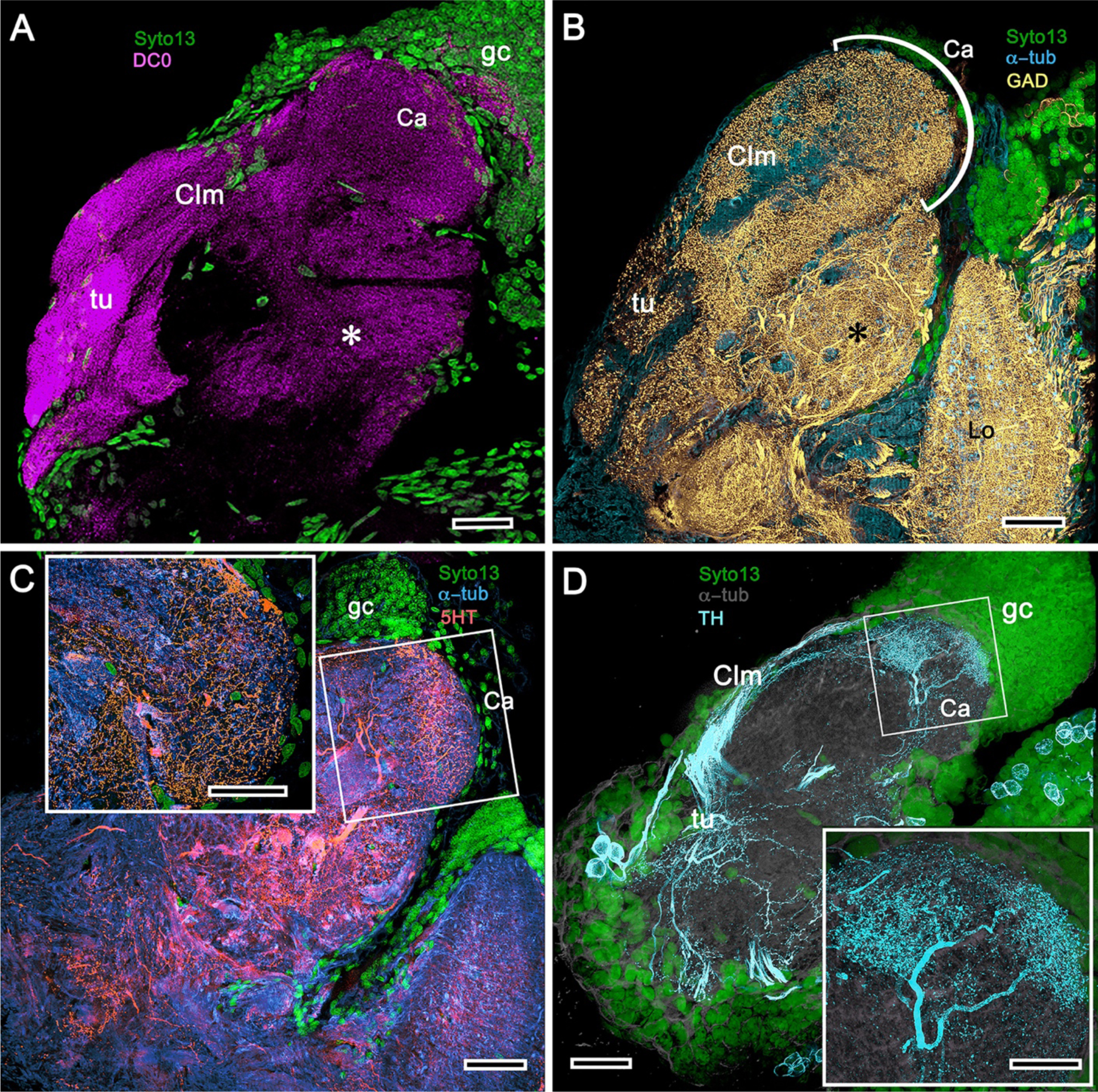
Mushroom body of *Crangon franciscorum*. **A.** Immunohistochemically labelled sections reveal moderate anti-DC0 expression in the mushroom body calyx (Ca) of *C. franciscorum* but a system of intensely labelled columnar lobes (Clm) ending as tubercular swellings (tu). As in carideans and reptantians, this species reveals lower levels of anti-DC0 immunoreactivity (asterisk) in the more distal parts of the lateral protocerebrum, beneath the calyx. **B.** Anti-GAD immunoreactivity is abundant in the calyx and the tubercles, but is sparser in the columnar lobes. Anti-GAD immunoreactivity also extends throughout the lateral protocerebrum and lobula (Lo). **C.** Anti-5HT labelling reveals putative serotoninergic processes in the calyx (inset: details shown in single scan) and regions within the lateral protocerebrum. **D.** Putative dopaminergic neurons, labelled with anti-TH (cyan), branch in two distinct territories of the calyx (Ca), enlarged in the inset lower right (gray, *α*-tubulin). Scale bars in A-D, 50μm, both insets, 20μm.

### Clade Reptantia

Reptantia are estimated to have originated in the mid-late Devonian (Porter et al., 2005; Wolfe et al., 2019). They are a natural group comprising genera that all possess a novel satellite neuropil in the deutocerebrum, adjacent to the olfactory lobe (Sandeman et al., 1993). This center, called the accessory lobe, comprises small spherical synaptic configurations that in some species number in the thousands. It receives connections from the olfactory lobes, but has no direct sensory input. Its projection neurons (PNs) contribute axons to the olfactory globular tract (OGT), which bifurcate at the same level as the central body to then extend collaterals to anti-DC0-positive centers in both lateral protocerebra.

### Transformed mushroom bodies in recent astacids

Fossil and geologically calibrated molecular phylogenies place the origin of the infraorder Astacidea, the lineage providing what are generally known as “lobsters” and “crayfish” at 250– 400 mya (Wolfe et al., 2019). Here we identify derivations of the ancestral mushroom body in the lateral protocerebra of the North American freshwater species *Orconectes immunis* (calico crayfish) and *Procambarus clarkii* (Louisiana crawfish).

Sullivan and Beltz’s 2001 description of the hemiellipsoid body of *Procambarus clarkii* identified its overall organization as two layered volumes, which we have named the cupola and torus (Strausfeld and Sayre, 2019). Neither volume appears outwardly distinct, the matrix of each comprising many microglomeruli (Figure 12A). However, closer inspection shows that these are larger and more densely packed in the torus than the cupola (Figure 12A, inset). Golgi impregnations, described in Strausfeld and Sayre (2019), resolve further differences between these two neuropils: each possesses a constrained organization of intrinsic neurons, which have their dendrites and short axons limited to either the cupola or torus. Both of these volumes also contain elaborate systems of local interneurons and glomerular ensembles, suggestive of calycal levels intermixed with local circuits, orthogonal arrangements appearing more pronounced in the torus. Thus, the cupola and torus each suggest a highly compressed modification of the mushroom body ground pattern. Anti-DC0 immunoreactivity shows both cupola and torus as usually, but not invariably, highly immunoreactive (compare Figure 12C with Figure 12D,E), as are discrete regions beneath it, here referred to as the subcalycal neuropil area VII (Figure 12C: nomenclature from Blaustein et al., 1988) and, laterally, the reniform body (RB, Figure 12C). Depending on the angle of sectioning, the outer level is domelike – hence the nomenclature cupola – whereas the inner level appears to have a toroidal architecture – hence torus. These volumes are matched by the distribution of anti-GAD immunoreactivity (Figure 12E). Organization of the cupola and torus as two independent units is also matched by the distribution of profuse anti-TH-immunoreactive efferent neurons and more sparsely distributed anti-5HT-immunoreactive processes (Figure 12F-H). Recordings from these neurons show them to be multimodal output neurons, comparable to MBONs of mushroom body columns (Mellon and Alones, 1997; McKinzie et al., 2003).

**Figure 12.**
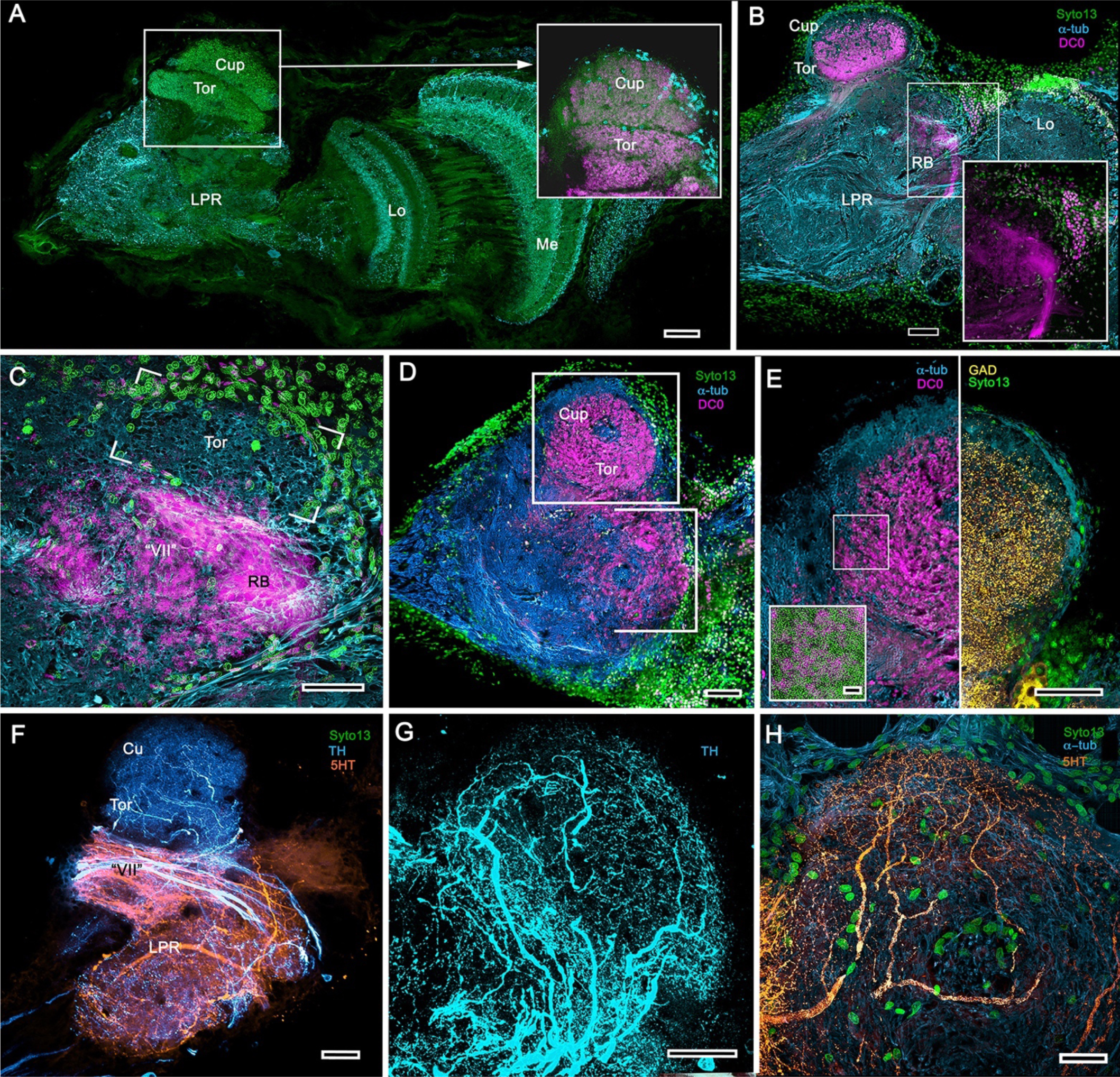
Transformed mushroom bodies in Astacidea. **A.** An overview of the crayfish lateral protocerebrum showing its distinctive hemiellipsoid body composed of two components, the rostral cupola (Cup) overlying the torus (Tor). All panels show the right lateral protocerebrum, where rostral is upwards and distal is to the right (f-actin, green; anti-allatostatin, cyan). Inset to A. anti-synapsin/f-actin immunostaining (magenta-green) shows the torus equipped with denser synaptic clusters than the cupola. However, it is the neuropil beneath the torus, receiving a massive input from the olfactory lobes that shows the greatest synaptic density. **B.** Anti-DC0 immunolabelling here shows the torus as more densely labelled. Neuropils ascribed to the reniform body (RB) are further distal, at the border between the lateral protocerebrum (LPR) and the optic lobe’s lobula (Lo). The inset shows a cluster of anti-DC0-positive cell bodies at the rostral surface associated with the reniform body like those identified in Brachyura (see Figure 15E,F). **C.** A section just glancing the torus, here showing almost no anti-DC0 immunoreactivity in the torus, but substantial anti-DC0 labelling in sub-calycal neuropils corresponding to neuropil (VII) described by Blaustein et al. (1988) and the probable location of the reniform body (RB). **D.** Intense anti-DC0 labelling of both levels of the hemiellipsoid body and neuropils beneath and distal to this center (bracketed). **E.** Alignment of the antiDC0-positive hemiellipsoid body in D with that of another specimen immunolabelled with anti-GAD. Discrete small glomerulus-like aggregates resolved by anti-DC0 (enlarged in inset) contrast with the uniform distribution of GAD-immunoreactive profiles. The inset demonstrates that anti-DC0-immunolabelled aggregates comprise anti-synapsin and f-actin labelled (magenta-green) synaptic microglomeruli comparable to those identified in the stomatopod mushroom body calyces (Wolff et al., 2017). **F-H.** Antibodies against anti-TH (cyan) and anti-5HT (orange) demonstrate distributions of these efferent neurons, which correspond to multimodal parasol cells (efferent neurons equivalent to MBONs) described by Mellon (2003) and McKinzie et al. (2003). Scale bars in A, B, D, F, 100μm; E, G, H, 50μm; inset to E, 2μm.

### Mushroom body hypotrophy of the ghost shrimp Neotrypaea californiensis

*Neotrypaea californiensis* is a species of ghost shrimp belonging to the 150-300mya (Triassic) infraorder Axiidea (Wolfe et al., 2019). This species inhabits shallow, intertidal waters covering muddy substrates in which it constructs deep burrows (often shared by individual *Betaeus harrimani*). Fossils of the subfamily Callianassinae, to which *Neotrypaea* belongs, first appear in the Eocene (56 mya; Hyžný and Klompmaker, 2016).

Labelling brains of the ghost shrimp *Neotrypaea californiensis* with anti-DC0 reveals paired mushroom body-like centers, which, because the species lacks eyestalks, are contained in foreshortened lateral protocerebra entirely within the head. As described in an earlier study (Wolff et al., 2012), where these centers were referred to as hemiellipsoid bodies, their neuropil is divided into two stacked volumes. The lower is intensely labelled by anti-DC0, whereas the upper volume, which comprises many hundreds of microglomerular synaptic sites, is not (Supplement to Figure 12). Thus, despite the current absence of immunostaining with other antibodies, this center has clear mushroom body attributes: a microglomerular calyx surmounting a condensed and foreshortened column, the neuropil of which is associated with a cluster of anti-DC0-immunoreactive perikarya in an adjacent globuli cell cluster.

### The chimeric mushroom body of Paguroidea and its evolved derivative

Originating in the Devonian, Reptantia gave rise to two branches, one providing the infraorders Achelata, Polychelida and Astacidea, the other providing the infraorders Axiidea and Gebiidea, and infraorders Anomura and Brachyura (together Meiura). Fossil-calibrated molecular phylogenies (Wolfe et al., 2019) demonstrate the superfamily Paguroidea within the infraorder Anomura diverging in the late Triassic to provide six families of which two, Paguroidae and Coenobitidae, are marine and land hermit crabs, respectively. Coenobitidae is the youngest, diverging approximately 35 million years ago. The earlier branching Paguroidae comprises marine hermit crabs. Here we compare anti-DC0-immunoreactive centers in the lateral protocerebrum of the marine hairy hermit crab *Pagurus hirsutiusculus* with those of the land hermit crab *Coenobita clypeatus* (McLaughlin et al. 2010).

In *Pagurus*, affinity to anti-DC0 (Figure 13A) reveals two immediately adjacent calycal domains in the lateral protocerebrum that together are continuous with an anti-DC0-positive columnar lobe. The neural architecture of the calyces (Figure 13B-D,F-H) is remarkably similar to the organization of afferents and intrinsic neurons in successive layers of the multistratified anti-DC0-immunoreactive hemiellipsoid body of the land hermit crab *Coenobita clypeatus* (Figure 13I-O). As demonstrated by Golgi impregnations (Wolff et al., 2012; Strausfeld and Sayre 2019), both *Pagurus* and *Coenobita* have corresponding networks provided by a variety of morphological types of intrinsic neurons. These form precisely defined orthogonal networks (Figure 13B,C) of dendritic processes and axon terminals. In *Pagurus*, one population of intrinsic neurons provides axon-like extensions into the lobes (Strausfeld and Sayre, 2019). In *Coenobita*, all of the intrinsic neuron axons are constrained to within the layered arrangements of its hemiellipsoid body (Wolff et al., 2012). The interpretation of the *Pagurus* center as representing a transitional organization is further supported by neurons corresponding to mushroom body output neurons (MBONs) already recognized in insect and stomatopod mushroom bodies. In *Pagurus*, the dendrites of anti-5HT- and anti-TH-immunoreactive MBONs intersect intrinsic neuron terminals in the tubercular swellings at the end of the columnar lobes. The layered arrangements of the dendrites of these immunoreactive neurons reflect the repeat organization of precisely aligned intrinsic neuron dendrites across each calyx’s width (Figure 13C,E).

**Figure 13.**
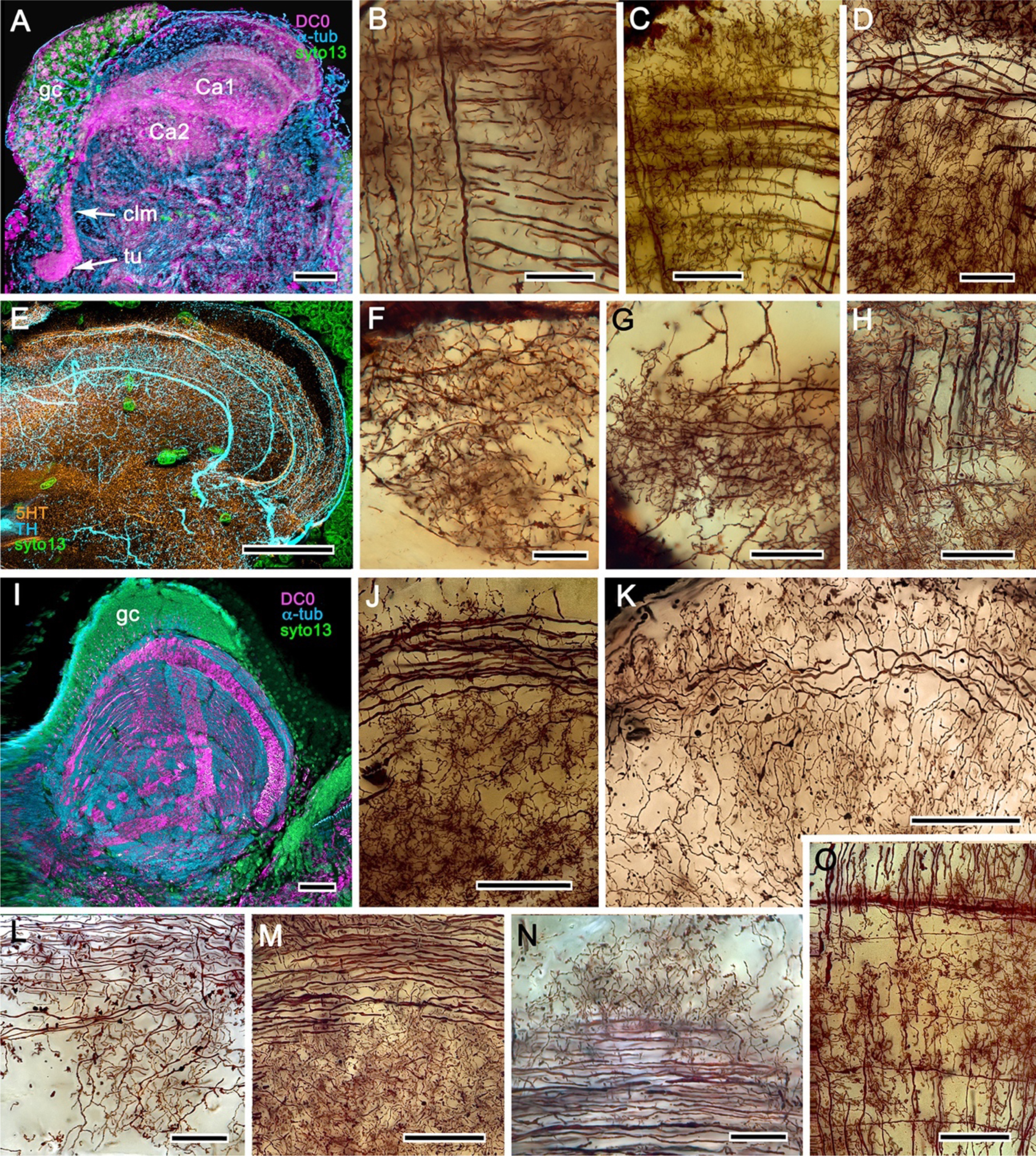
The reptantian mushroom body with and without its columnar lobe. Golgi impregnations demonstrate corresponding orthogonal arrangements in the two calyces of *Pagurus* (Ca1, Ca2 in panel **A**), which extend as a columnar lobe (clm) ending in tubercular swellings (tu), and in the multistratified hemiellipsoid body of *Coenobita* (**I**), which lacks a lobe. **B-D, F-H**. Golgi impregnations of *Pagurus*’s calyces show characteristic orthogonal (B,F,G) and rectilinear arrangements (C) of intrinsic neurons invested by extended beaded fibers from the olfactory globular tract (OGT; F) and providing axon-like processes that extend to the lobe (D; see also, Strausfeld and Sayre, 2019). Anti-5HT and anti-TH immunostaining resolves corresponding efferent dendritic trees aligned with orthogonal networks (E). These are shown for the multistratified *Coenobita* hemiellipsoid body (panel **I**). **J-O**. Golgi impregnations of the *Coenobita* hemiellipsoid body demonstrate orthogonal and rectilinear organization of its intrinsic neurons. Scale bars in A, 50μm; B-D, 25 μm; E, 50μ; F, 10μm; G, 25 μm; H,J,K, 50 μm; I, 100μm; L-O, 25 μm.

### The hemiellipsoidal mushroom body of *Coenobita clypeatus*

The anomuran family Coenobitidae, to which the Caribbean land hermit crab belongs, is a recent group appearing approximately 35-40 mya (Bracken-Grissom et al., 2014a; Wolfe et al., 2019). *Coenobita clypeatus* and its cousin the coconut crab *Birgus latro* have attracted considerable attention regarding olfactory adaptations associated with terrestrialization (Stensmyr et al., 2005). However, the juveniles develop in a marine habitat and, as expected, their antennular olfactory receptor neurons are typical of marine species in possessing ionotropic receptors, the axons of which terminate in large deutocerebral olfactory lobes (Harzsch and Hansson, 2008; Groh et al., 2014).

As shown in both *B. latro* and *C. clypeatus,* massive relays from the olfactory lobes project to the lateral protocerebra to terminate in greatly inflated centers referred to as hemiellipsoid bodies (Harzsch and Hansson, 2008). Subsequent studies showed these neuropils to be defined by numerous DC0-positive stratifications (Figure 14A,B). Golgi impregnations and electron microscopy demonstrated that these tiered arrangements consist of orthogonal networks of intrinsic neurons, postsynaptic to afferent terminals and presynaptic to dendrites, resulting in an organization that corresponds to circuitry recognized in the insect mushroom body’s columnar lobe (Brown and Wolff, 2012; Wolff et al., 2012). Volumes of neuropil situated laterally beneath the dome are also anti-DC0positive, as is a globular center corresponding to the reniform body (Figure 14A,B). The correspondence of this center’s internal organization with that of a mushroom body’s column is further demonstrated by the arrangements of dendritic trees belonging to output neurons, which are equivalent to MBONs. Figure 14C shows that anti-TH- and anti-5HT-immunoreactive dendrites extend across strata, each of which is also defined by synaptic configurations revealed by f-actin and anti-synapsin (Figure 14D) and, in Figure 14F, by anti-*α*-tubulin and an antibody against PKA-RII that regulates DC0 (see Methods). Antibodies against GAD reveal broadly distributed processes that are not aligned within rows but extend across them (Figure 14C, E).

**Figure 14.**
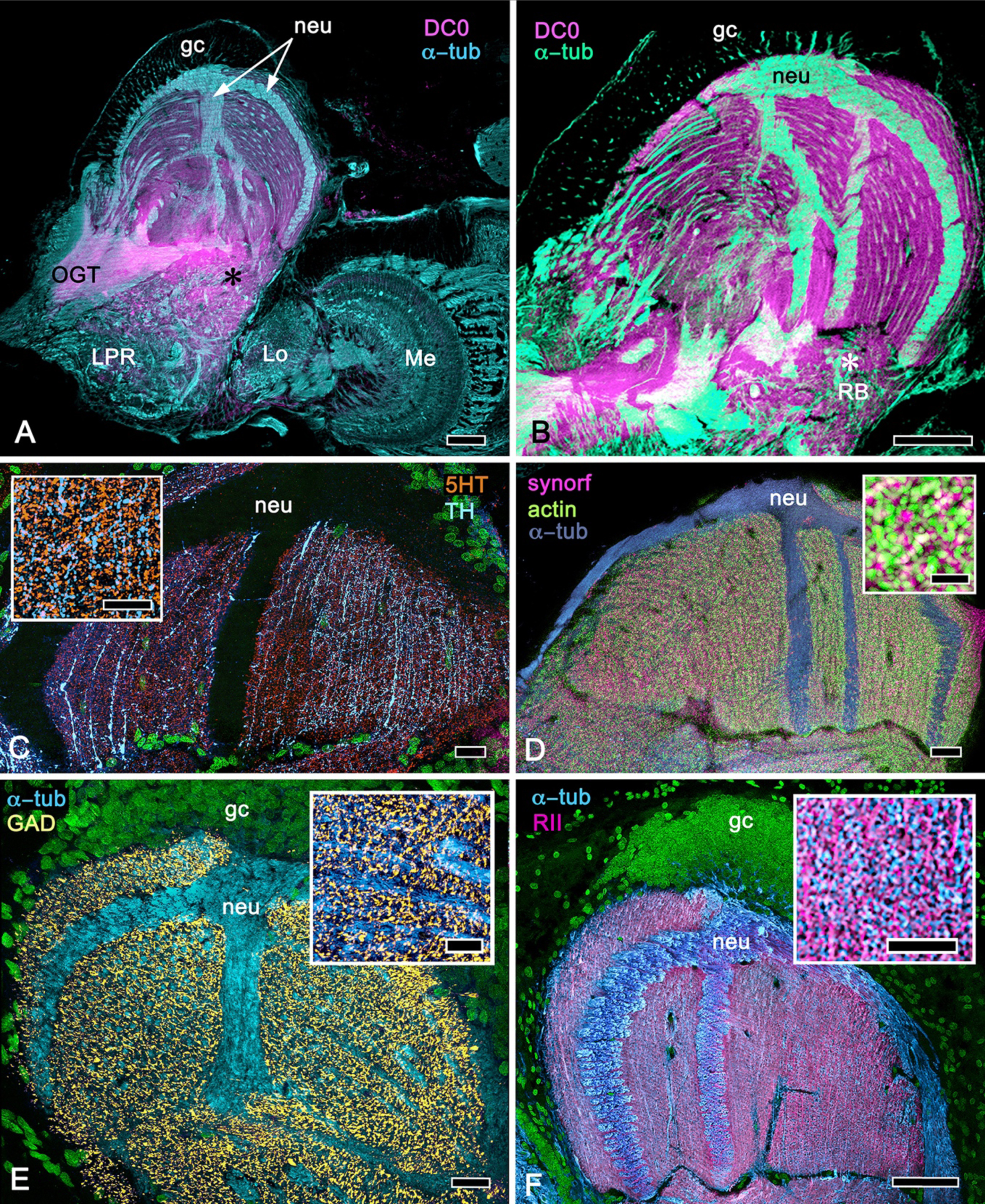
Calycal hypertrophy in the *Coenobita clypeatus* mushroom body. **A,B.** DC0 and other immunological markers reveal a characteristic system of nested strata comprising orthogonal fibers that originate from the intrinsic neurons (neu; cyan, anti-*α*-tubulin). Anti-DC0 (magenta) defines these strata as well as the entrance of the olfactory globular tract (OGT) into the calyx and several regions of the lateral protocerebrum beneath the calyx, including the reniform body (RB, asterisk in A,B). Other regions of the lateral protocerebrum (LPR,) and optic lobe (Lo, lobula; Me, medulla) show little or no affinity. **C.** 5HT-(orange) and TH-immunoreactive (cyan) fibers in the calyx extend within each of the DC0-positive strata but are notably absent from the neurite bundles of intrinsic neurons. The distribution of TH is not uniform (inset), but patchy indicating discrete domains of TH processes within the stratified intrinsic neuron networks. **D.** Anti-synapsin (magenta) and f-actin (green) demonstrate regions of dense synaptic connectivity indicated by microglomerular configurations (inset). In this panel, intrinsic neuron bundles labelled with anti-*α*-tubulin (blue). **E.** Distributed processes labelled with antisera against GAD extend across strata, labelling neuropil occupied by afferents to the calyces (inset). **F.** RII, an antiserum developed in *Drosophila melanogaster* against the regulatory subunit of PKA, confirms an expression pattern corresponding to that of DC0 (**A,B**) and is shown comingling with neuropil in and between stratifications (inset). Scale bars in A,B, 100μm; C-E, 20μm; F, 50μm; inset C, 5μm; inset D, 2μm; insets E-F, 5μm.

This stratified organization of the anomuran mushroom body/hemiellipsoid body is probably older than suggested by Paguridae or Coenobitidae. It is also revealed by silver stains of the lateral protocerebrum of the squat lobster *Munida quadrispina*, a member of the Galatheoidea. This superfamily has been indicated as arising in the Triassic-Jurassic (Bracken-Grissom et al., 2014a; Porter et al., 2005).

### Extreme divergence of the mushroom bodies in Brachyura

Brachyura (true crabs) is a species-rich infraorder, now comprising about 6,800 species (Ng et al., 2008), that diverged 240 mya (Wolfe et al., 2019). Here we consider two species, both of which spend intertidal periods out of water; the shore crab *Hemigrapsus nudus* (Varunidae) and fiddler crab *Uca* (*Minuca*) *minax* (Ocypodidae). Fossil-calibrated molecular phylogenies indicate a mid to late Cretaceous origin for the Varunidae and Ocypodidae (Tsang et al., 2014; Wolfe et al., 2019).

The opportunistic detritovore *H. nudus* lives in a flat visual ecology interrupted by occasional pebbles and rocks. These shore crabs often come into immediate contact with each other, where they may initiate contact-induced actions, such as defensive aggression involving jousting and pushing, but there is no obvious ritualized courtship as there is in fiddler crabs. Individuals employ crevices as transitory residences, responding to predatory threats by opportunistically retreating and hiding in them (Jacoby, 1981). It is likely that they can deploy allocentric memory, judging from the habit of near- or fully-grown individuals to retreat to a specific location when threatened. These actions contrast with those of fiddler crabs, such as *Uca mina*x – deposit feeders living on mudflats – that are renowned for their ritualized courtship displays and rapid path integration to established home sites to escape predation (Crane, 1975), and visual behaviors that relate to compound eyes adapted for vision in a flat world (Zeil and Hemmi, 2006).

There are few recent studies on the brains of crabs, exceptions being descriptions of the optic lobes of the South American shore crab *Neohelice granulata* (Sztarker et al., 2005), a description of the crab’s reniform body (Thoen et al., 2019) and a comparative study on the brains of fully terrestrial crabs (Krieger et al., 2015). However, as yet no clearly defined center, comparable to that of an achelate or anomuran “hemiellipsoid body,” has been convincingly identified, possibly due to difficulties of antibody penetrance in the lateral protocerebra by anti-FMRFamide and anti-synapsin, the antibodies used for screening in the Krieger et al., 2015 account. This in part explains past imprecision in defining the location of the “hemiellipsoid body” and why it is conflated with the “medulla interna” (Krieger et al., 2012), and difficulties in identifying an appropriate candidate. For example, a description of NMDA receptor distribution in *Neohelice granulata* diagrammatically indicates a “hemiellipsoid body” occupying about a quarter of the caudal volume of the lateral protocerebrum (Hepp et al., 2013), a position opposite to that shown for any other crustacean and opposite to that shown by antibodies raised against DC0 and RII.

Treatment of eyestalks of six *H. nudus* with those antibodies resulted in four of twelve eyestalks showing barely detectable immunoreactivity with one specimen exhibiting some immunoreactivity in a tributary of the OGT. Eight consistently showed various levels of immunoreactivity in regions of the rostral lateral protocerebrum. Figure 15A-F shows three of five specimens that all resolved strong immunoreactivity of large parts of the rostral neuropil associated with clusters of globuli cells lying over the lateral protocerebrum’s rostral surface. One cluster of globuli cells also supplies a welldefined anti-DC0-immunoreactive neuropil positioned immediately proximal to the lobula. This matches the elaborations of the reniform body (Fig.15E), as resolved in Golgi impregnations and Bodian reduced silver-stained specimens (Thoen et al., 2019). This center is known to participate in visual habituation and was understandably (but mistakenly) interpreted by Maza et al. (2016) as a mushroom body. However, as shown by Thoen et al. (2019), this center is not a homologue of the mushroom body or the hemiellipsoid body. Depending on the tilt of sectioning and the section’s depth the intensely anti-DC0-immunoreactive pedestal of the reniform body is resolved extending obliquely from the rostro-dorsal to the caudal margin of the lateral protocerebrum (Figure 15A-F). Sections of fortuitously oriented blocks can reveal the entire reniform body complex, here shown digitally enhanced against other anti-DC0 territories in Figure 15E,F.

**Figure 15.**
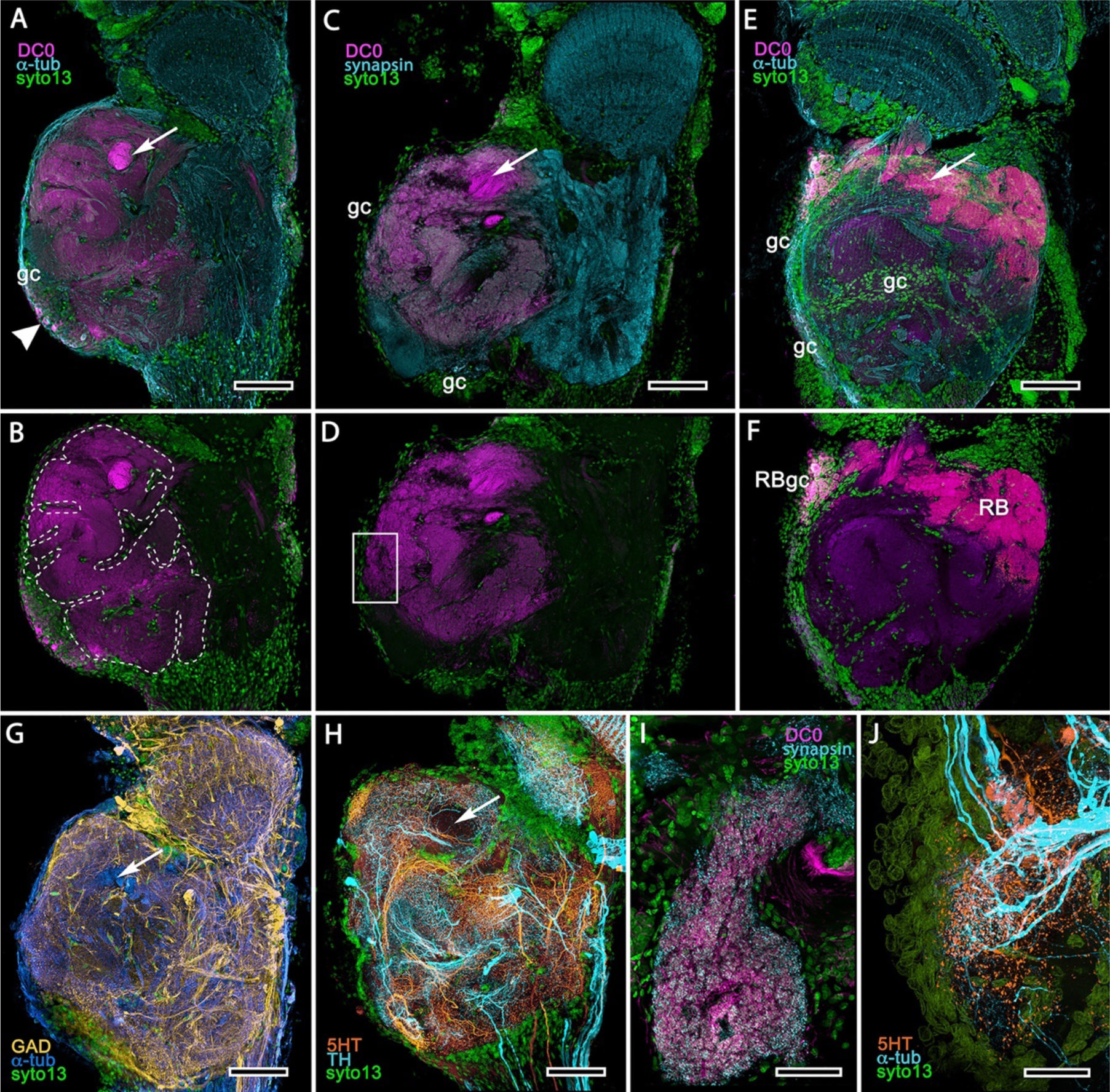
Mushroom body homologues of the shore crab *Hemigrapsus nudus*. **A-H** depict sections of the right lateral protocerebrum: rostral surface is to the left; distal (towards the optic lobes) is the top of each panel. **A-F.** Anti-DC0-immunoreactive territories are interpreted as modified mushroom bodies, lacking columnar lobes and enormously expanded in the rostral lateral protocerebrum. Distally, each of these centers is penetrated by the pedestal (bright magenta in A-F, arrow in panels A-H) belonging to the large reniform body typifying varunid crabs. The entire territory occupied by the reniform body (RB) is shown in E,F against the dimmed magenta surround. Anti-DC0 labelling is shown at normal intensities in panels A-D. The maximum extent of the mushroom body neuropil is indicated by the dotted outline in B. Cell bodies of intrinsic neurons (syto13, gc) occur scattered over the rostral and dorsal surface of the lateral protocerebrum (A-F) some also showing elevated anti-DC0 immunoreactivity (e.g., arrowhead in A). Globuli cells providing the reniform body (RBgc) also show anti-DC0 immunoreactivity, as indicated in panel F. **G.** Distribution of anti-GAD-immunoreactive (yellow) processes mainly in caudal volumes of the lateral protocerebrum and the lobula. **H.** Distribution of anti-5HT(orange) and anti-TH-immunoreactive (cyan) processes. **I,J.** Labelling with anti-DC0 (magenta) and the synaptic marker SYNORF1 (cyan; I) and with anti-5HT (orange) and anti-TH (J) reveal discrete regions (one boxed in D) in anti-DC0-immunoreactive territories that are suggestive of mushroom body-like circuitry. Scale bars in A-H, 100μm; I, J, 25 μm.

Figure 15 depicts territories of the lateral protocerebrum of *H. nudus* indicating that almost the entire rostral half of the neuropil is occupied by what is likely a highly modified mushroom body. These very large anti-DC0-immunoreactive domains, characterized by their cortex-like boundary of fissures and lobes (Figure 15B), are reminiscent of the arachnid mushroom body of Amblypygidae, which is also extensively folded (Wolff and Strausfeld, 2015). In *H. grapsus*, the volume associated with the mushroom body is relatively free of anti-GAD immunoreactivity, whereas antibodies raised against 5HT and TH suggest that anti-TH may be more abundant in the lateral protocerebrum’s upper half than in the lower. The fiddler crab, *Uca minax* (Supplement to Figure 15), reveals similar anti-DC0-immunoreactive territories that also appear to be highly folded. Less extensive than those in *H. nudus*, the two most immunoreactive territories are separated by a volume showing lower affinity to anti-DC0. Notably, vibratome sections treated with SYNORF1 (anti-synapsin) and f-actin to resolve synaptic microglomeruli, selectively reveal only specific subterritories of the volume occupied by the anti-DC0-positive center (inset: Supplement to Figure 15).

Scrutiny of anti-DC0 territories identifies anti-synapsin as well as anti-TH and anti-5HT immunoreactive arrangements suggestive of mushroom body-like configurations (Figure 15I, J), the locations of which correspond to the boxed area of Figure 15D. Current analysis of this obviously elaborate and challenging system is currently underway, focusing on Golgi impregnation to identify the distribution of OGT afferents and interneurons associated with them, with the aim of isolating arrangements that suggest mushroom body-like circuits or other circuit configurations that may be unique to Brachyura.

## DISCUSSION

Here we demonstrate that antibodies raised against DC0 identify discrete integrative centers in the pancrustacean lateral protocerebrum that point to divergent evolution from the mushroom body ground pattern (Figure 16). Antisera against GAD, 5HT and TH resolve neurons associated with these centers that are consistent with the immunohistological signatures of inhibitory circuits and input and output neurons that characterize insect mushroom bodies. Golgi impregnations demonstrate that populations of small basophilic neuronal cell bodies, called globuli cells, provide the center’s intrinsic neurons. This neurohistological palette identifies evolved mushroom body transformations: simplification, loss, and elaboration to the dome-like architectures, historically referred to as hemiellipsoid bodies.

**Figure 16.**
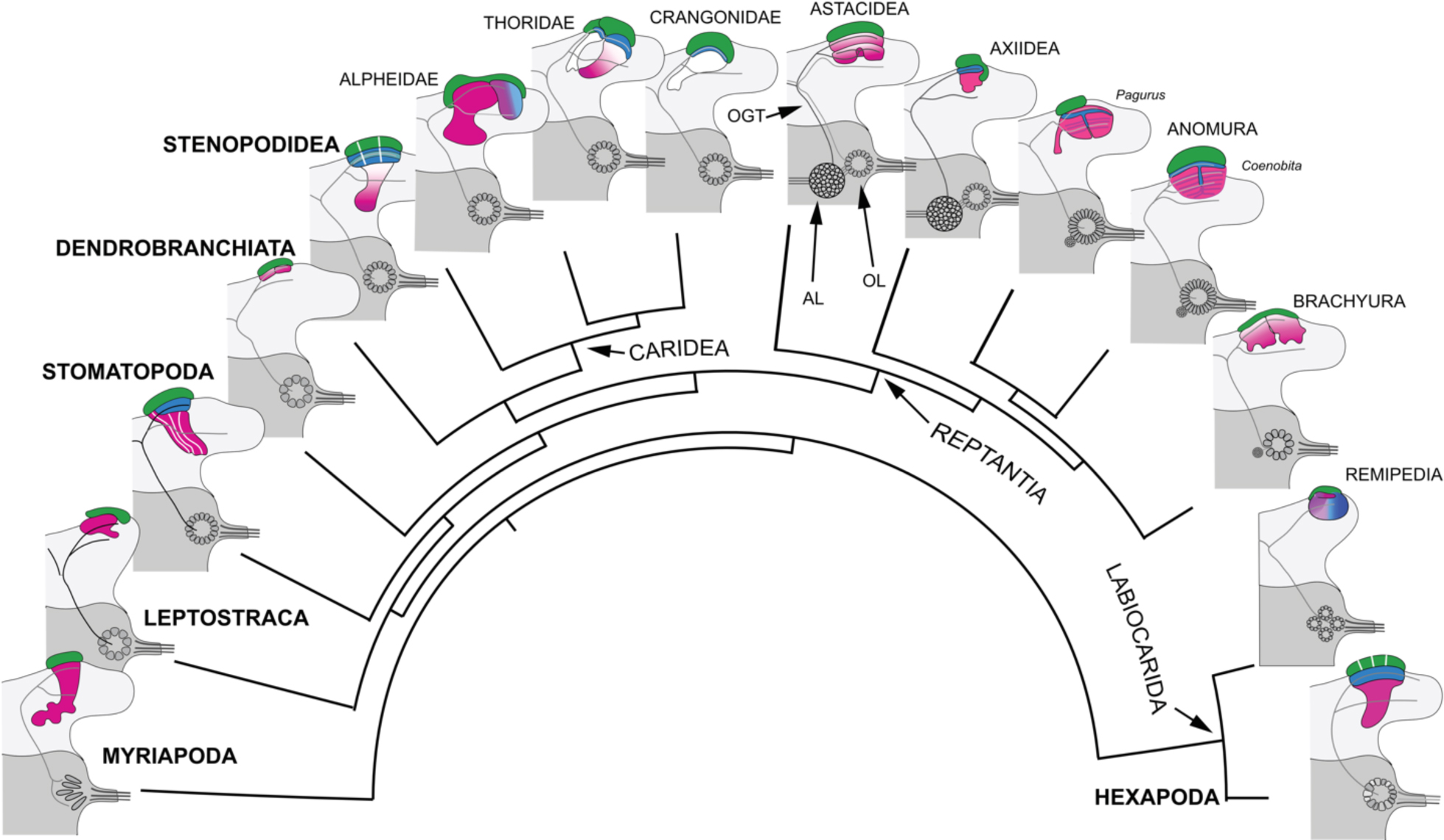
Retention and divergence of the mushroom body ground pattern in Mandibulata. Schematized shapes of mushroom bodies described in this study (for actual profiles see Figures 1, 17) and their evolved derivatives are mapped onto a pancrustacean molecular phylogeny (after: Oakley et al, 2013; Wolfe et al. 2019; Schwentner et al., 2017), here extended to include the mandibulate outgroup Myriapoda (represented by the chilopod mushroom body; Wolff and Strausfeld, 2015). Each schematic depicts the right lateral protocerebrum without its optic lobes. Hemisegments of the deutocerebrum and protocerebrum are indicated by dark and light grey shading, respectively. The deutocerebrum is shown with its olfactory lobe (OL); or, in Reptantia, the olfactory and adjoining accessory lobe (AL). The antennal glomerular tracts (OGT) are shown with their contralateral extension indicated in all examples except Chilopoda and Hexapoda, where the OGT is exclusively homolateral. Magenta indicates anti-DC0/RII identification of corresponding neuropils; green indicates globuli cell clusters; blue indicates distinct calycal organization. Schematics for Caridea are, clockwise, Alpheidae, Thoridae, and Crangonoidea. Anomura are represented by *Pagurus* and *Coenobita*. Evolved diminution is here shown for Dendrobranchiata. Despite miniaturization in Leptostraca, the RII-immunoreactive center reveals reduced columnar lobes. A comparable arrangement is resolved in Remipedia (see Fig. 3 in: Stemme et al., 2016).

### Transformation of the mushroom body to the astacid/achelate “hemiellipsoid” morphology

Decapods are traditionally grouped into two clades, “Natantia” (natare, to swim) and “Reptantia” (reptare, to creep), rationalized as swimming decapods being morphologically distinct from those that walk. Natantia is a paraphyletic group embracing the infraorders Dendrobranchiata, Stenopodidea, and Caridea (Scholtz and Richter, 1995). Stenopodidea and many Caridea are, as adults, agile reef dwellers negotiating complex topographies; it is these taxa that possess centers closest to the mushroom body ground pattern.

Most descriptions have, however, focused on Reptantia (e.g., Derby and Blaustein, 1988; Wachowiak and Ache, 1994; Mellon and Alones, 1997; Mellon, 2003; McKinzie et al., 2003; Schmidt and Mellon, 2010); morphological and molecular phylogenetics demonstrate this group as monophyletic (Scholtz and Richter, 1995; Porter et al., 2005; Wolfe et al., 2019). Fossil-calibrated molecular phylogenies estimate reptantian time of origin at 400–385 mya (Wolfe et al., 2019; Porter et al., 2005).

Reptantia are unified by possessing a unique apomorphy called the accessary lobe (Hanström, 1925; Sandeman and Scholtz, 1993; Sandeman et al., 1995). In contrast to an olfactory lobe comprising turbinate or cuneate units, traditionally (but inappropriately) called “glomeruli,” the spherical or approximately crescent-shaped accessory lobes are composed of a mass of diminutive spherules usually arranged as an outer shell enclosing an inner volume (Sandeman and Scholtz, 1993). The spherules receive inputs exclusively from the adjacent olfactory lobe (Sandeman and Luff, 1973; Wachowiak et al., 1996). Whereas in natantians the olfactory lobe provides relays to the mushroom bodies, in the reptantian infraorders Achelata, Astacidea, and Axiidea it is the accessory lobe that provides the inputs (Sandeman and Scholtz, 1993; Sullivan and Beltz, 2001, 2004, 2005), with axons from the olfactory lobes supplying underlying lateral protocerebral neuropils (Sullivan and Beltz, 2001, 2004, 2005).

As shown here, and in Strausfeld and Sayre (2019), although not equipped with columnar lobes the achelate hemiellipsoid body consists of two distinct neuropils reflecting an evolved transformation from the ancestral mushroom body ground pattern. Internal organization, comprising elaborate networks of shortaxoned and anaxonal intrinsic neurons, is provided by globuli cells further supporting mushroom body ancestry. Efferent neurons (parasol cells) shown from intracellular recordings to relay multisensory associations from these centers have been proposed to correspond to mushroom body output neurons (McKinzie et al., 2003; Mellon et al., 1992); like MBONs, these neurons from hemiellipsoid bodies are resolved using antisera against TH and 5HT (Figure 12F-H).

In the reptantian infraorders Anomura and Brachyura accessory lobes are inconspicuous. It is the olfactory lobes that supply the welldefined mushroom body of the anomuran *Pagurus* (Figure 13A-H) and its distinctively layered homologue in Coenobitidae (Figures 13I-O,14). Organization in Brachyura is far more difficult to interpret. Anti-DC0 immunocytology of the shore crab *Hemigrapsus nudus* reveals a prominent immunoreactive domain, denoted by fissures and lobes, that occupies almost the entire rostral half of the lateral protocerebrum (Figure 15) proximal to the reniform body (Thoen et al., 2019). Two territories similarly occupy the lateral protocerebrum of the fiddler crab *Uca minax* (Supplement to Figure 15). These observations do not align with a study of terrestrial crabs based on synapsin immunocytology, indicating hemiellipsoid bodies with achelate-like substructures (Krieger et al., 2015). But synapsin indiscriminately denotes any synaptic neuropil, and delineations offered in that account preclude comparisons. Further, the designation of these domains in crabs as “hemiellipsoid bodies” (Krieger et al., 2015) would suggest that they have derived from the hemiellipsoid body morphology. There is no evidence yet to support this, particularly when the anti-DC0-immunoreactive regions shown here suggest a folded neuropil, reminiscent of the mushroom bodies of whip scorpions belonging to the aranean order Amblypygi (Wolff and Strausfeld, 2015).

How fast was the change from the ancestral mushroom body to an achelate-astacidtype hemiellipsoid morphology? Was this change unique to Reptantia? Did the hemiellipsoid body morphology evolve in concert with the accessory lobes? Might this have occurred before reptantian diversification in the late Devonian? Evidence not limited to Reptantia suggests that such changes could have been quite rapid. In the Crangonidae, *Crangon franciscorum* has a mushroom body-like calyx and columnar lobes, whereas its sister species *Paracrangon echinata* has a bulbous hemiellipsoid body-like center and a diminutive lobe (Figures 10C,D,11). *Crangon* and *Paracrangon*, which occupy different habitats (see below), are estimated to have diverged in the early Eocene, about 56 mya (Davis et al., 2018).

### Evolved reductions and losses

Reduction and loss are possible explanations for very small or absent centers corresponding to the mushroom body/hemiellipsoid bodies. Such second-order olfactory integration centers may not be required in species that live in conditions where associative memories are of negligible relevance. The minute anti-DC0-immunoreactive centers of *Penaeus*, shown in Figure 3A, suggest such an evolved reduction in a migratory species that lives some of its adult life in the featureless ecology of the off-shore water column. In certain isopods, possibly olfactory specialists, the mushroom body is reduced to a mere vestige, as shown in Figure 3B. In terrestrial isopods, a complete loss is suggested to relate to diminution of the olfactory pathway (Harzsch et al., 2011). Complete loss of the center also is reported for species constrained to unusual ecologies, such as isolated fresh water caverns (Ramm and Scholtz, 2017; Stegner et al., 2015). The marine isopod *Saduria entomon* may have a greatly reduced hemiellipsoid body (Kenning and Harzsch, 2013), but this may be insecure as well because unspecific antibodies cannot properly delineate neuropils. As cautioned by Ramm and Scholtz (2017), because the olfactory globular tract (OGT) terminates in the much reduced lateral protocerebra of isopods (Stemme et al., 2014) this does not imply the presence of a hemiellipsoid body.

Leptostraca is sister to all Eumalacostraca (Wolfe et al., 2016). Until now, evidence for a center comparable to a mushroom body has been tenuous. An absence of this center would be problematic because if mushroom bodies and their modifications are a defining feature of Pancrustacea, then mushroom bodies would be expected to occur in this basal malacostracan group. Neuroanatomy of the *Nebalia* brain (Figure 17A-C) reveals a small lateral protocerebrum crowned by large eyes, supplying at least three nested optic lobe neuropils (Figure 17C; also, Sinakevitch et al., 2003). One diagnostic trait of Multicrustacea is that the OGT links each olfactory lobe with both lateral protocerebra. In the leptostracan *Nebalia herbstii* this bifurcation is present but much reduced (Kenning et al., 2013) and we have observed the same in *N. pugettensis*.

**Figure 17.**
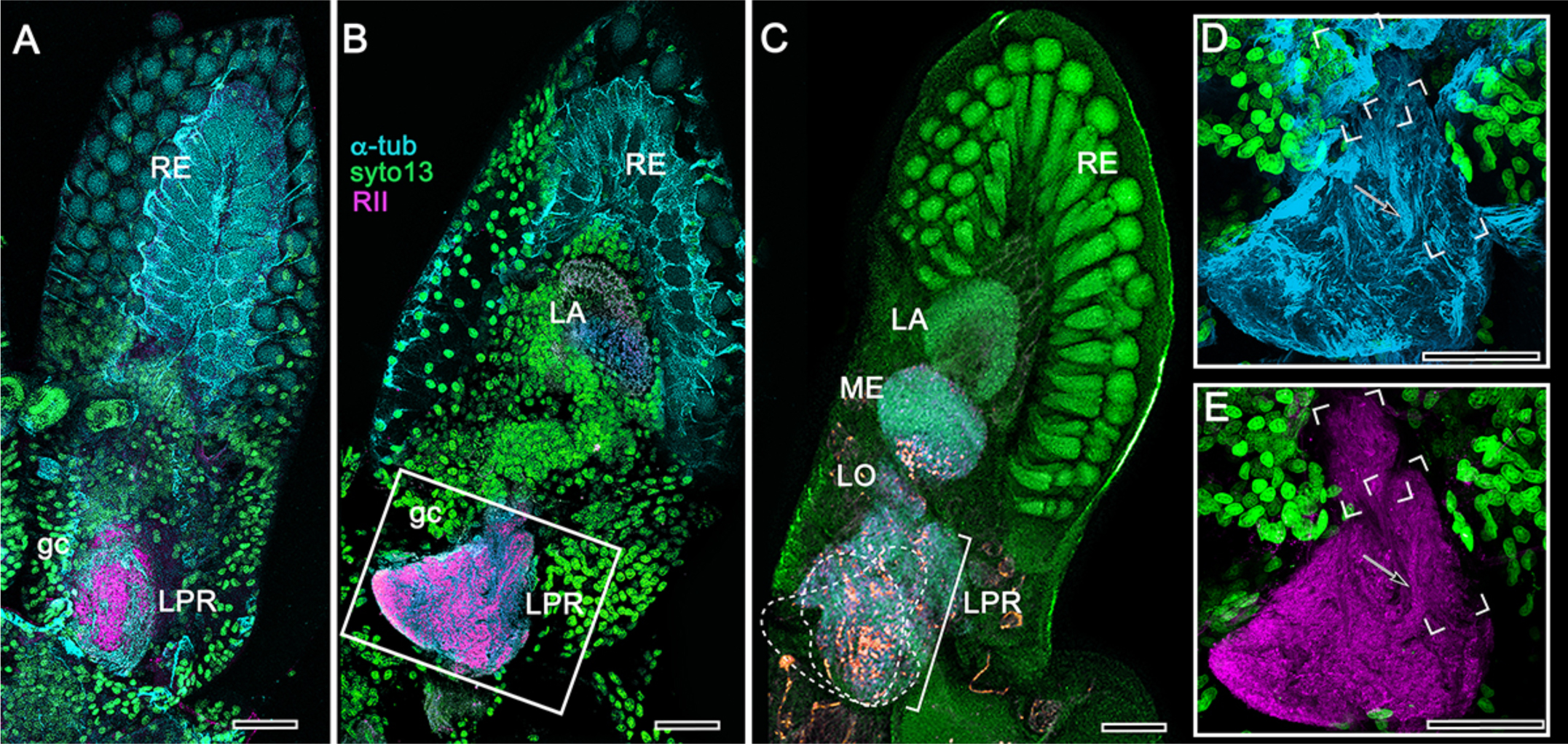
Mushroom body homologues in *Nebalia pugettensis.* Two successive sections shown in panels A, B, reveal an ensemble of neuropils strongly immunoreactive to antibodies raised against RII (see Methods). An anti-RII immunoreactive region of neuropil (A), expanding into a planar calyx (B) gives rise to two small but well-defined columnar lobes extending forwards. These domains are superimposed in panel C, beneath which are deeper levels of the lateral protocerebrum (LPR) comprising several discrete neuropils central to the nested optic lobe neuropils (LA, lamina; ME, medulla; LO, lobula). The boxed area in panel B is enlarged in D, E. Panel D shows the neuroarchitecture of the calyx resolved with anti-*α*-tubulin; the columnar lobes are framed, one showing bundled parallel fibers (arrow). E. shows these as anti-RII labelled processes (also arrowed in E) extending from the triangular calyx. RE, retina; gc, globuli cells. Orange in panel C is FMRFamide-immunoreactivity. Scale bar indicates 50μm.

Kenning et al. (2013) were unable to demonstrate a clearly defined hemiellipsoid body in *N. herbstii* and wrote that it was “ambiguous,” but nevertheless attributed its location to the “medulla internis.” Our application of anti-RII to the brain of *Nebalia pugettensis* identifies an anti-RII immunoreactive neuropil that comprises a relatively large shallow cap over the rostro-dorsal lateral protocerebrum (Figure 17A, B). The cap provides two very short columns, in which anti-D-tubulin and anti-RII immunoreactivity suggest parallel arrangements of fibers (Figure 17D, E). However, this observation is from a single specimen, albeit revealing these structures in both lateral protocerebra. Small perikarya, suggestive of globuli cells, are contiguous with a larger population also supplying optic lobe neuropils (Figure 17A, B). The smallness of the mushroom body in *Nebalia* suggests a possible evolved miniaturization, comparable to that described from hexapod Collembola (Kollmann et al., 2011).

Although the central complex in *Nebalia* is as prominent as it is in decapods (Kenning et al., 2013), miniaturization of the mushroom body may not be surprising if the simple morphology (reflecting behavior) of *Nebalia* is compared with *Cascolus ravitis,* the oldest known fossil stem representative of this group (approximately 430 mya: Siveter et al., 2017). Since the mid-Silurian, leptostracans have undergone morphological simplification (Dahl, 1985) and extant species (e.g., Martin et al., 1996) are mostly epibenthic scavengers or suspension feeders limited to burrowing habits in simple ecologies (McCormack et al., 2016).

Exceptions to the expectation that life in the water column might drive evolved reduction are recognized in two groups, both of which lack compound eyes. In certain predatory Copepoda, small intertidal members of Multicrustacea, glomerular antennular lobes are connected by heterolaterally projecting OGTs to prominent second-order olfactory centers located in their lateral protocerebra. These centers are, for the size of the brain, large. They consist of synaptic networks originating from numerous minute globuli cell bodies (Andrew et al., 2012). This arrangement conforms to the malacostracan ground pattern, as does that of an allotriocarid lineage Remipedia, also a class of predatory crustaceans.

Molecular phylogeny identifies Remipedia as the closest crustacean relative of Hexapoda (Oakley et al., 2013; Schwentner et al., 2017). Yet each of its olfactory lobes sends axons that bifurcate to anti-DC0-immunoreactive centers in both lateral protocerebra (Stemme et al., 2016). Each center gives rise to a small columnar-like extension (Stemme et al., 2016), also visible in thin sections as a short stubby column extending from the larger neuropil (see Fig. 6A, B in Fanenbruck and Harzsch, 2005). This is reminiscent of the arrangement in *Nebalia* (Figure 17B, D, E). Indeed, a conundrum that still requires resolution is that overall brain organization in Remipedia is much closer to that of Malacostraca than it is to its molecularly related allotriocarids (Fanenbruck et al. 2004). One of those is Hexapoda; another is Cephalocarida, the most basal member of Allotriocarida (Schwentner et al., 2017). In both, mushroom bodies are connected homolaterally to their olfactory lobes (Stegner and Richter, 2011), contrasting with the bilateral connections in Remipedia and Malacostraca.

### Divergent evolution of the pancrustacean mushroom body: correlations with ecologies and behaviors

What behavioral attributes unite pancrustacean species that possess columnar mushroom bodies? How might these differ in those that possess the hemiellipsoid body morphotype? Here we speculate that the aerial-terrestrial ecology exploited by insects is comparable to elaborate three-dimensional ecologies of reefs and escarpments exploited by stomatopod and caridean species. Making allowance for what is a miniscule sample size, our results suggest that highly mobile species occupying dynamic three-dimensional ecologies have retained the mushroom body ground pattern, which in insects provides continuous updating of spatial associations and their valences (Cognigni et al., 2018). That mushroom bodies are crucial for spatial awareness is supported by studies demonstrating that they increase in size as a response to acquiring information about threedimensional space (Kühn-Bühlmann and Wehner, 2006; Montgomery et al., 2016; van Dijk et al., 2017). Anomurans such as land hermit crabs, in which circuits characterizing mushroom body lobes have been subsumed into the calyces, have refined spatial memory for exploration and homing (Krieger et al., 2012). Crustaceans and insects possessing columnar mushroom bodies negotiate structurally elaborate three-dimensional habitats, or actively hunt within defined territories. Ancestral mandibulates likely evolved in comparable ecologies. Micro-CT studies of their appendicular morphologies suggest that the oldest pancrustaceans from the lower Cambrian were locomotory adepts rather than simple animals that crawled on the sea bed (Zhai et al., 2019).

Life on the ocean floor, intertidal flats, or the bed of a stream or lake, such as that adopted by the reptantians Achelata, Astacidea, and Axiidea, may require altogether different computational networks for coping with what is predominantly a planar world. Such distinctions are also suggested by recent Crangonidae as discussed above. Large columnar mushroom bodies typify highly active species, such as those that show place memory. Examples are cleaner shrimps such as *Stenopus hispidus*, motile reef dwellers such as *Lebbeus groenlandicus* or *Spirontocaris lamellicornis*, and active hunters such as Stomatopoda. Mushroom bodies are enormous in pistol shrimps – the only crustacean group that includes eusocial species – as they are in their cousin *Betaeus*, a genus commensal with burrowing Axiidea and required to memorize location. Carideans, such as *Palaemonetes pugio* that negotiate the complex geometries of aquatic plants, are similarly equipped with prominent mushroom body-like centers.

### Evolutionary diversity of crustacean mushroom bodies, but stability of the insect homologue, may also relate to convergent evolution of their olfactory supply

The primary sensory input to mandibulate mushroom bodies is olfactory, and the olfactory lobes of crustaceans and insects have been promoted as homologous (Schachtner et al., 2005; Harzsch and Krieger, 2018). However, the present study demonstrates that the mushroom body ground pattern has undergone profound morphological, and it must be assumed, functional divergence across malacostracans (Figure 16). In contrast, as shown from a comparable study of the insect mushroom body (Strausfeld et al., 2009), there has been far-reaching morphological conservation of the mushroom body ground pattern across dicondylic insects.

Up to the level of the protocerebrum, scrutiny of the crustacean and insect olfactory pathways suggests that these share surprisingly little in common, and may be very different indeed. If those differences, which are discussed next, relate to different modes of sampling and encoding of odorant space, might organization typifying the crustacean olfactory pathway have permitted relaxed evolution of the mushroom body, whereas organization typifying the insect olfactory pathway constrained divergent mushroom body evolution?

Distinctions between the crustacean and insect olfactory systems begin at the level of the uniramous appendages of the insect and crustacean deutocerebrum. Although they are segmental homologues across Mandibulata (Boxshall, 2004), the morphology of the crustacean “antennule” diverges from that of the insect “antenna,” which due to the loss of the second antenna in that group assumes some of the second antenna’s functionality. But the first crucial distinction applies to the morphologies of olfactory sensilla and their olfactory receptor neurons (ORNs). Aesthetascs, the flexible odorant sensilla of crustaceans, are distributed in malacostracans on the antennule’s lateral flagellum. Each sensillum can contain some hundreds of ORNs, the dendrites of which may further branch to provide extensive membrane surfaces (Tierney et al., 1986; Grünert and Ache, 1988; Schmidt and Mellon, 2010; Hallberg and Skog, 2010). In insects, four principal morphologies of odorant sensillum can contain 1-6 poorly branched or unbranched ORNs (Schneider and Steinbrecht, 1968; Altner and Prillinger, 1980). An exception is the hymenopteran placode sensillum serving 30+ ORNs (Getz and Akers, 1994). These receptor-level distinctions are amplified by differences of sensory transduction mechanisms. Crustacean ORNs are exclusively equipped with ionotropic channels (Derby et al., 2016). In dicondylic insects, in addition to ionotropic ORNs, the majority of ORNs employ liganded-gated channels, there being as many variants of these in a species as there are genes that encode them (Robertson et al., 2018).

Differences between insects and crustaceans are profound in the olfactory lobes, which in crustaceans comprise identical wedge- or lozenge-shaped synaptic glomeruli. ORN axons terminate in one or several of these (Tuchina et al., 2015), but there is no hard evidence that any specific type of ORN targets a specific subset of glomeruli. The opposite applies to insects. Generally, each glomerulus has a specific location (and size and shape) and is functionally unique in receiving converging terminals from, usually, a single genetically identical set of ligand-gated ORNs (Fishilevich and Vosshall, 2005), there being as many glomeruli as there are genes encoding ligand-gated receptors (Couto et al., 2005). These arrangements comprise an odortypic map: a feature not present in crustaceans.

Mushroom bodies are supplied by relay neurons (projection neurons: PNs) originating from the olfactory lobes. This feature is entirely different in crustaceans and insects. Hundreds – in some species thousands – of axons belonging to PNs extend rostrally from the crustacean olfactory lobe (or in certain reptantians from the accessory lobes) to reach the lateral protocerebrum, many axons dividing into two tributaries extending to both sides. In crustaceans, no PN confines its dendrites to a single unit of the olfactory lobe: PNs have wide-field dendrites extending to numerous glomeruli (Wachowiak and Ache, 1994: Schmidt, 2016). This sharply contrasts with insects where each glomerulus contains the dendritic trees of 3-5 PNs, and only a small population of PNs is associated with groups of glomeruli. Thus, relatively few PN axons extend rostrally from the insect antennal lobe and their axons exclusively target the ipsilateral protocerebrum to end in the mushroom body and the distally adjacent neuropils of the lateral horn (Galizia and Rössler, 2010). The only crustacean with comparable homolateral PN projections is Cephalocarida (Stegner and Richter, 2011), an allotriocaridid related to Hexapoda (Oakley et al., 2013).

Thus, up to the lateral protocerebrum, each level of the olfactory pathway typifies either a crustacean or an insect (Table 2). No crustacean possesses ligand-gated ORNs, and hexapod molecular phylogenetics implies that the insect odorant receptor family (ORs; Brand et al., 2018) appeared after terrestrialization (Missbach et al., 2014). In crustaceans, arrangements amongst ORN terminals and widely branching PNs implies all-to-all connectivity. There is no evidence for functionally related differentiation of olfactory glomeruli, and it is currently assumed that reconstruction of chemical and temporal patterning of the odorant milieu is distinct from its encoding by the insect system (Derby and Schmidt, 2017). Could these differences explain why insect mushroom bodies are all similar and why crustacean ones are not? It is these fundamental differences that need to be addressed in asking whether perhaps the labelled-line organization of the insect olfactory pathway has constrained the evolution of the mushroom bodies in insects, whereas the all-to-all crustacean system has allowed divergent adaptations of computational circuits, each ecologically matched at the level of the mushroom bodies or their evolved derivatives.

### The “medulla terminalis” does not exist

The entrenched opinion is that central to its optic lobe neuropils, the crustacean lateral protocerebrum comprises an integrative center called the “medulla terminalis,” with which the hemiellipsoid body is invariably associated (Krieger et al., 2012). The term is an artificial construct for a volume of the lateral protocerebrum that corresponds vaguely to the collective neuropils comprising an insect’s lateral protocerebrum (Strausfeld, 1976, 2019; Ito et al., 2014). The term “Medulla terminalis” was introduced in the 1890s, to indicate a (non-existent) fourth optic neuropil, and later adopted in 1925 by Hanström to indicate a volume of neuropil little understood at the time. The term should be dropped as it trivializes both the complexity of the eumalacostracan lateral protocerebrum and has the effect of permitting vague indications of the presence of a “hemiellipsoid body” when the actual delineation of such a neuropil cannot be resolved, as in the 2019 description by Wittfoth et al. Proposing equivalence of the “medulla terminalis” with the insect’s lateral horn (see, Harzsch and Krieger, 2018), which is a distal complex supplied by the antennal lobe and optic glomeruli (Galizia and Rössler, 2010; Ito et al., 2011; Strausfeld et al., 2007) also detracts from deeper scrutiny and evolutionary discussion. Thus far, the study by Blaustein et al. (1988) is the only serious attempt to give names to definable volumes in the crustacean lateral protocerebrum other than the mushroom bodies and their evolved derivatives.

### Mushroom bodies unify Mandibulata (Pancrustacea and Myriapoda)

Even as Bellonci (1882) identified and named the mantis shrimp’s corpo emielissoidale, he stated that with its columnar extension – the corpo allungato – the two components were identical to the insect mushroom body. It is also a leitmotiv of Hanström’s crustacean studies that he refers to these centers as corpora pedunculata, irrespective of whether they are pedunculate (Hanström, 1925, 1931). In his extensive 1931 study of decapod brains, Hanström stated throughout that hemiellipsoid bodies are mushroom bodies that either lack the columnar lobe or have incorporated it. Yet, in the ensuing years the belief that the domed hemiellipsoid bodies of crustaceans are fundamentally different structures from the lobed mushroom bodies of insects has become not only widely accepted, but specifically advocated as a basis for the ground pattern of the malacostracan brain (Sandeman et al., 2014; Wittfoth et al., 2019; Machon et al., 2019). The doctrine has become entrenched: that hemiellipsoid bodies are a derived trait of crustaceans, whereas mushroom bodies are a derived trait of hexapods (Sandeman et al., 2014; Machon et al., 2019).

That it is difficult to relinquish the hemiellipsoid body as the ground pattern is exemplified by a recent paper referring to the stomatopod mushroom body’s calyx as a hemiellipsoid body but categorically declining any consideration of its columnar lobes (Machon et al., 2019). Shoe-horning mushroom bodies into a hemiellipsoid morphology, as in descriptions of the caridean *Rimicaris exoculata* (Machon et al., 2019; loc. cit. Fig. 12), stifles any discussion about evolution, as does the assertion of the hemiellipsoid body’s ground pattern status, even in species where the neuropil is not demonstrable (Wittfoth et al., 2019). The recent demonstration of mushroom bodies in the Stomatopoda and other lineages described here present evidence that the mushroom bodies provide the ground pattern for the crustacean protocerebral olfactory center. At the very least, this demonstration necessitates a reconsideration of the relationship between the mushroom body and structures referred to using Hanström’s (1931) synonym for the calyces. One of the goals of the present paper has been to show how the transformation from a columnar lobed mushroom body morphotype to a its calycal (hemiellipsoid) morphotype can be followed across eumalacostracan evolution (Figures 1B, 16) while unifying this center in Eumalacostraca and Allotriocarida. And, as we know from observations of Myriapoda (Diplopoda + Chilopoda), this mandibulate outgroup of Pancrustacea is likewise defined by its paired DC0-immunoreactive mushroom bodies (Wolff and Strausfeld, 2015, 2016).

It is indisputable that the variety of higher centers in crustaceans referred to as mushroom bodies or “hemiellipsoid bodies” contrasts with the uniformity of mushroom bodies across insects. To be sure, it is remarkable that no member of Dicondylia has been identified with an evolutionarily modified mushroom body comparable to a hemiellipsoid body. The single monocondylic order Archaeognatha has no mushroom body at all (Strausfeld, 2012), its phylogenetic position within Hexapoda (Giribet et al., 2004) implying an evolved loss.

The one fascinating exception is an experimentally induced point mutation that transforms the wild-type mushroom body to a hemiellipsoid body-like center. This is in the *Drosophila* brain mutant ks63 (mbd “deranged”), the paired centers of which lack columnar lobes because their Kenyon cell axons are constrained to within the voluminous hybrid (calyx + lobe) neuropil, supplied by normal projection neurons from the olfactory (antennal) lobes. The mutant flies are able to discriminate odors and show odor-induced behavior, but were not shown to be capable of learning odors (Heisenberg, 1980; Heisenberg et al., 1985).

That the mushroom body ground pattern has persisted with far less variation across Hexapoda than it has across Crustacea is no reason to dispute that it unites Hexapoda and Crustacea. Evidence provided here demonstrates that in Crustacea mushroom bodies are – to paraphrase Richard Owen’s famous statement (1843) – the same center under every variety of form and function. The notion that hemiellipsoid bodies are something phyletically separate and distinct is based on an historical misapprehension that diverts attention from why such a plentitude of variations evolved and what drove their evolution.

## ACKNOWLEDGEMENTS

The research described here is supported by the National Science Foundation under Grants No. 1754798 awarded to NJS and No. 1754610 to GHW, as well as the University of Arizona’s Center for Insect Science, and funding to NJS from the University of Arizona Regents’ Fund. Our gratitude is directed to Daniel Kalderon, Columbia University, New York, for supplying the DC0 and RII antibodies. We thank the staff of the University of Washington’s Friday Harbor Marine Laboratories, San Juan, for their hospitality and unfailing help in obtaining living specimens. We have much profited from exchanges with Joanna M. Wolfe, Harvard University, Cambridge, MA, who gave crucial advice on the use of time lines and identified a number of errors in our original assessments. We are indebted to Camilla Strausfeld for critically discussing versions of the final manuscript, suggesting many improvements and meticulously editing the text.

## SUPPLEMENTAL FIGURES

**Supplement to Figure 1.**
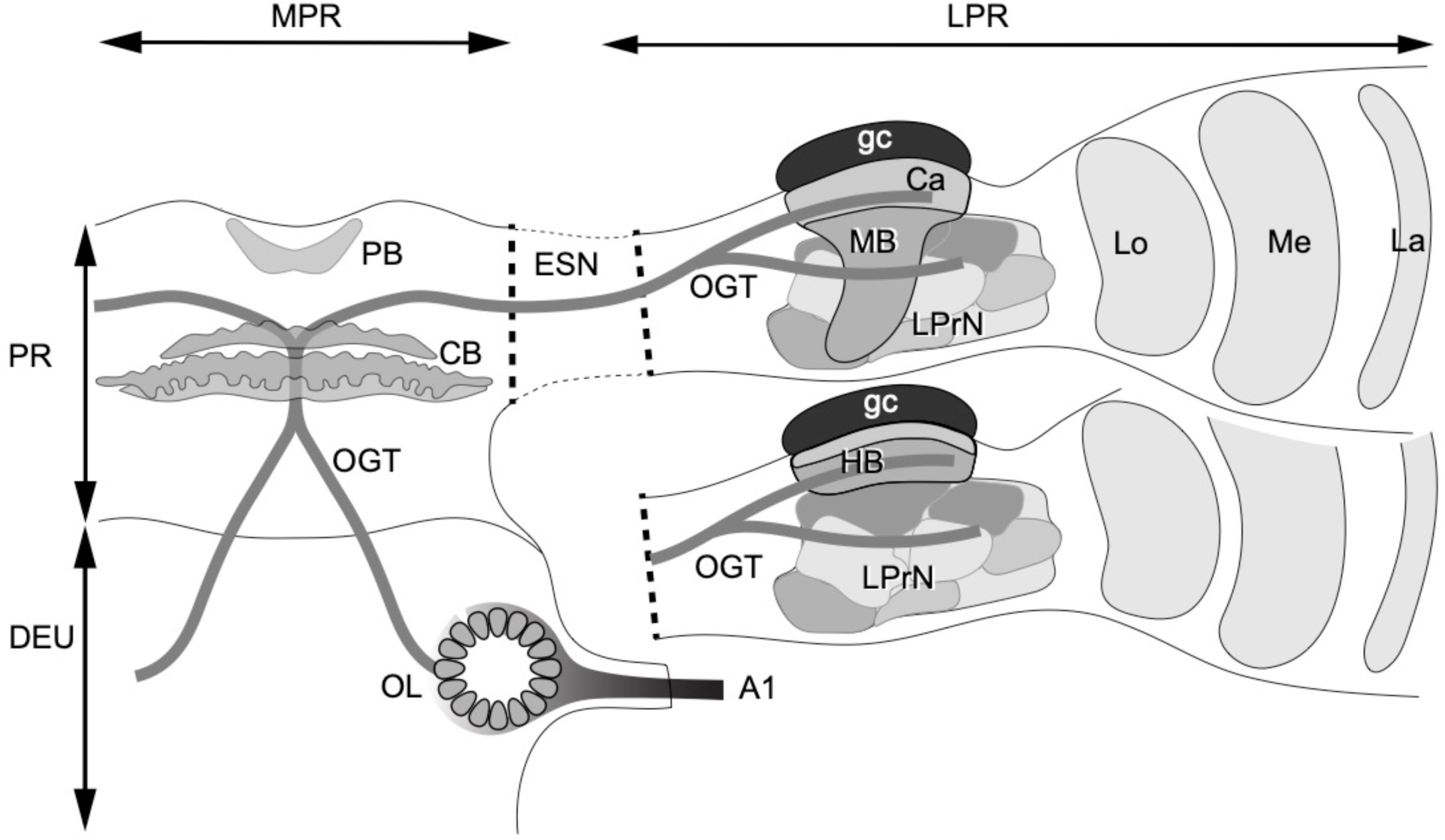
Ground pattern of the pancrustacean olfactory pathway and its lateral protocerebral termini. Olfactory receptor neurons supply the olfactory lobe (OL) through antennular nerve A1. Relays from the olfactory lobe (or, in certain Reptantia, the accessory lobe) are carried by the olfactory globular tract (OGT). This branches at the midline immediately dorsal to the central body (CB). Its ipsi- and contralateral axon branches extend to the lateral protocerebrum via the eyestalk nerve (ESN). Branches of the OGT supply the calyx (Ca) and several discrete lateral protocerebral neuropils beneath it (LPrN). An evolved loss of the mushroom body’s hemiellipsoid body (HB), as shown in lower eyestalk. Intrinsic neurons of the mushroom body or hemiellipsoid body originate from a dense population of neuronal perikarya called globuli cells (gc). Other abbreviations: MPr medial protocerebrum; Pr protocerebrum; Deu deutocerebrum; PB protocerebral bridge; La, Me, Lo first, second and third optic neuropils.

**Supplement to Figure 12.**
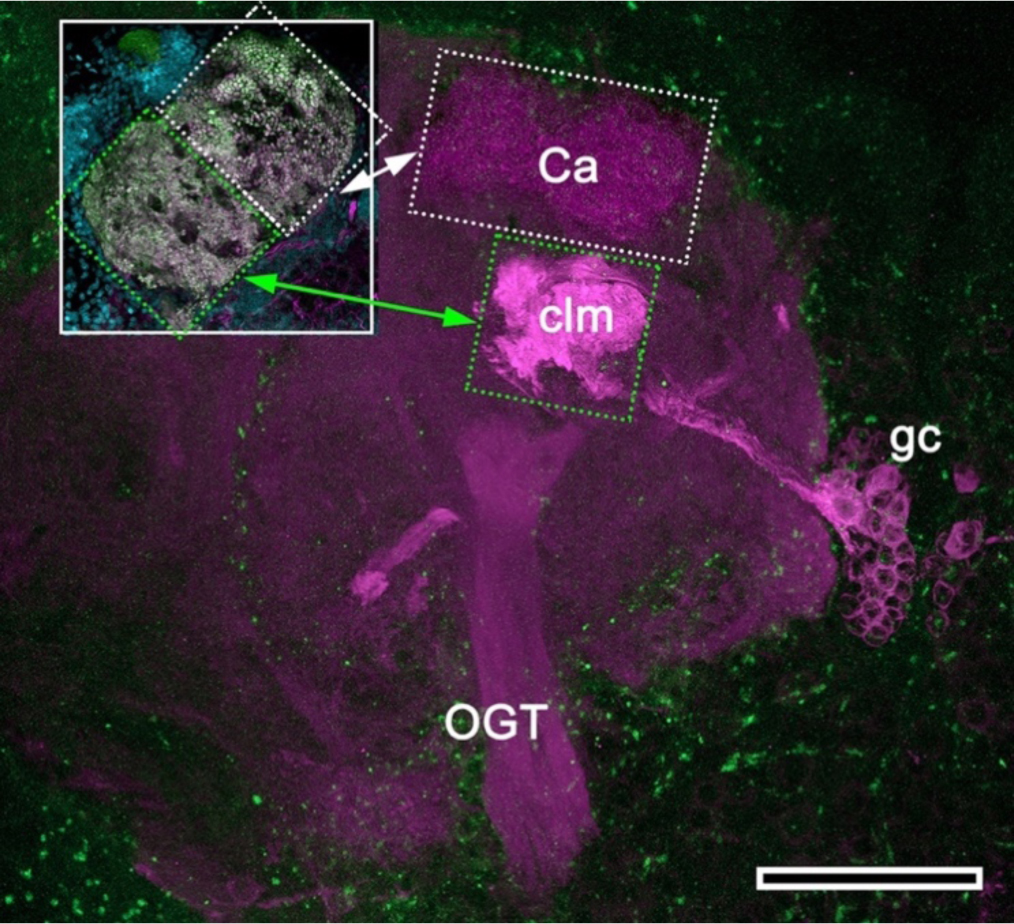
Mushroom body hypotrophy of the ghost shrimp *Neotrypaea californiensis*. (Modified from Brown and Wolff, 2012). Immunolabelled sections stained with antisera against DC0 (magenta) and phalloidin conjugated to Alexa 488 (green). Intrinsic cell bodies of the calyx (Ca) and elsewhere have not been stained in this preparation. However, a cluster of anti-DC0-immunoreactive intrinsic neurons (gc) gives rise to a greatly condensed column (clm) extending from beneath the calyx (Ca). The inset shows the calyx consisting of a dense population of microglomerular synaptic sites (inset; synapsin, magenta; f-actin, green; syto13, cyan). Very low anti-DC0 immunoreactivity typically shows the surrounding protocerebral neuropils and olfactory globular tract (OGT). Scale bar, 100μm.

**Supplement to Figure 15.**
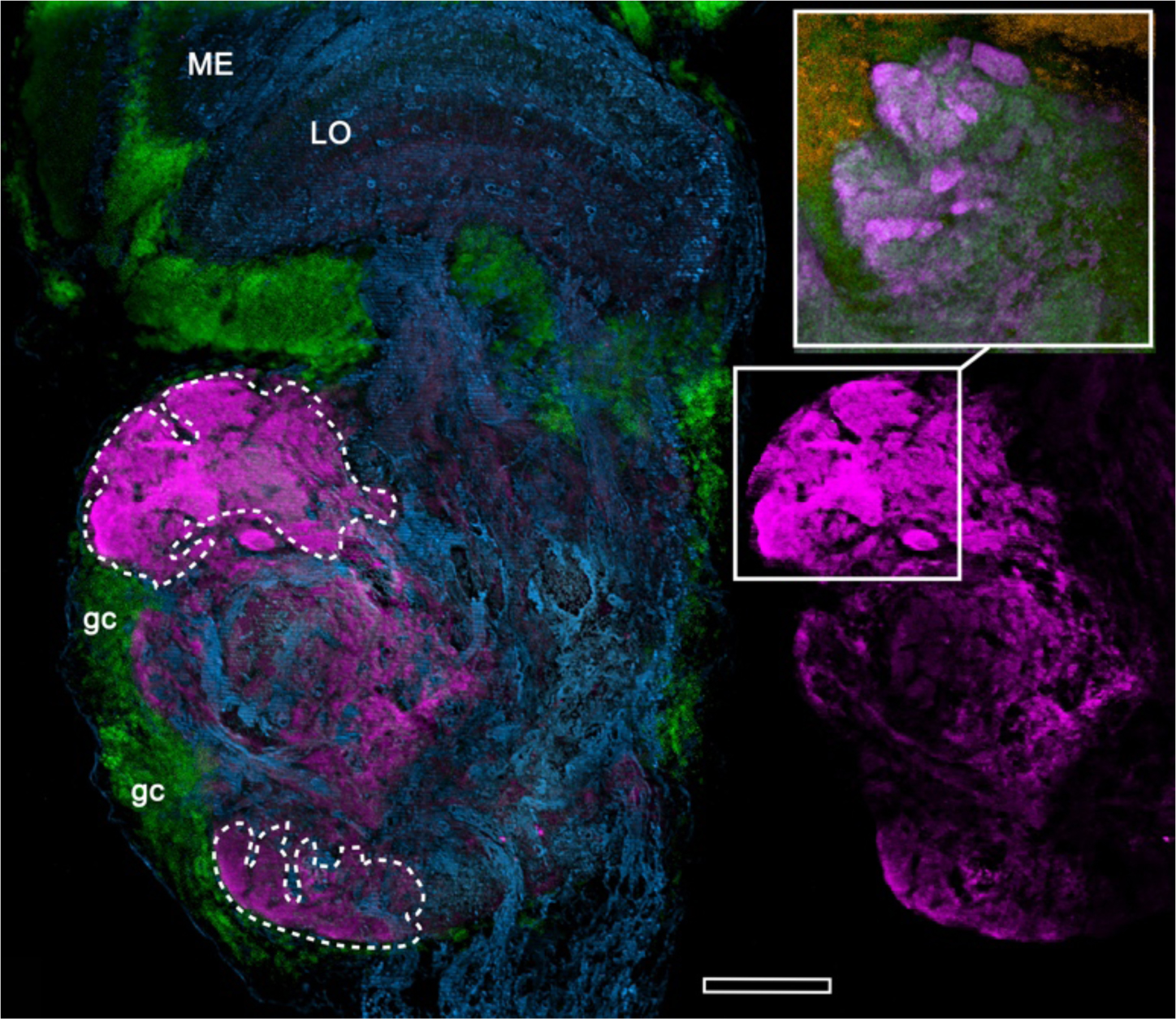
Anti-DC0-immunoreactive territories (magenta, lower right) of the fiddler crab *Uca minax* (rostral to the left, distal upwards). The left panel delineates the boundaries (dashed lines) of two of them, of which the most distal has dense immunoreactive folds and fissures. Staining with anti-synapsin (magenta) and f-actin (green, upper right inset; cell bodies, yellow) demonstrates the dense population of synaptic sites in the folds and fissures. ME, medulla; LO, lobula; gc, globuli cells. Scale bar is 100 μm.

